# Sexual coordination in a whole-brain map of prairie vole pair bonding

**DOI:** 10.1101/2023.07.26.550685

**Authors:** Morgan L. Gustison, Rodrigo Muñoz-Castañeda, Pavel Osten, Steven M. Phelps

## Abstract

Sexual bonds are central to the social lives of many species, including humans, and monogamous prairie voles have become the predominant model for investigating such attachments. We developed an automated whole-brain mapping pipeline to identify brain circuits underlying pair-bonding behavior. We identified bonding-related c-Fos induction in 68 brain regions clustered in seven major brain-wide neuronal circuits. These circuits include known regulators of bonding, such as the bed nucleus of the stria terminalis, paraventricular hypothalamus, ventral pallidum, and prefrontal cortex. They also include brain regions previously unknown to shape bonding, such as ventromedial hypothalamus, medial preoptic area and the medial amygdala, but that play essential roles in bonding-relevant processes, such as sexual behavior, social reward and territorial aggression. Contrary to some hypotheses, we found that circuits active during mating and bonding were largely sexually monomorphic. Moreover, c-Fos induction across regions was strikingly consistent between members of a pair, with activity best predicted by rates of ejaculation. A novel cluster of regions centered in the amygdala remained coordinated after bonds had formed, suggesting novel substrates for bond maintenance. Our tools and results provide an unprecedented resource for elucidating the networks that translate sexual experience into an enduring bond.

## Introduction

Bonds are essential to the social lives of many species, enabling individuals to coordinate parental care, territorial defense, or other shared activities (Emlen & Oring, 1977; Marlowe, 2000; Trivers, 1972). Among humans, friendships and a happy marriage protect against a wide range of stressors and their sequelae (Baumeister & Leary, 1995; Hawkley & Cacioppo, 2010), while social integration is one of the strongest predictors of reduced morbidity and mortality risk (Snyder-Mackler et al., 2020). Recent evidence suggests social buffering against stress-related disease can be found in many other mammals as well, including species of nonhuman primates, rodents, ungulates, and hyrax (Snyder-Mackler et al., 2020). Nonhuman animals offer unique insights into the mechanisms of social attachment and their consequences.

The neurobiology of bonding is most studied in the prairie vole, a socially monogamous rodent in which males and females form bonds, share a nest, and raise young together (Lieberwirth & Wang, 2014; Young & Wang, 2004). In prairie voles, as in many pair-bonding species, bonds form in response to courtship and repeated mating (Carter et al., 1995; Getz et al., 1993); indeed, copulation itself is regarded as a form of courtship that allows mutual assessment and coordinates reproduction (Dewsbury, 1988; Eberhard, 1996). Because of these characteristics, prairie voles have become an excellent model species to study neural circuits that underpin affiliative behaviors (Kenkel et al., 2021). Classic studies emphasized the roles of neuropeptides in reward circuitry (Aragona et al., 2006; Young & Wang, 2004), work that has been refined to reveal that neural ensembles in the nucleus accumbens encode approach behavior between mates (Scribner et al., 2020), and the strength of functional connections between nucleus accumbens and prefrontal cortex predicts social contact (Amadei et al., 2017). Mated prairie voles exhibit empathy-like consolation behavior that reduces markers of distress in both members of a pair (Burkett et al., 2016). In one common neuroanatomical model, some 18 different brain regions shape aspects of memory, reward and approach that are essential to bond formation and its consequences (Walum & Young, 2018). Despite this extraordinary work, we still have an incomplete understanding of the circuits that enable the experience of mating to become an enduring bond. Building upon this foundation will require a shift from a piece-meal to systems-wide perspective.

In several species, including wasps, fish, bats, mice, and humans, researchers using a variety of biological measures find that social interactions are accompanied by coordinated neural states (Hasson et al., 2012; Kingsbury et al., 2019; Kinreich et al., 2017; Long et al., 2020; Vu et al., 2020; Zhang & Yartsev, 2019). In fighting fish (*Betta splendens*) and paper wasps (*Polistes fuscatus*), shared brain-transcriptomic signatures occur in individuals immediately after competitive interactions (Uy et al., 2021; Vu et al., 2020). In Egyptian fruit bats (*Rousettus aegyptiacus*) and lab mice, freely interacting individuals have correlated neural activity in the frontal cortex, and increases in inter-brain correlations predict whether subsequent interactions occur (Kingsbury et al., 2019; Zhang & Yartsev, 2019). In humans, inter-brain EEG recordings become synchronized in a variety of interactive contexts (Hasson et al., 2012), such as during social gaze and positive affect in couples (Kinreich et al., 2017). Such data have led to the inference that closely interacting dyads, including bonded individuals, coordinate aspects of their physiological and neural states (Long et al., 2020).

In contrast to the emphasis on shared brain states, sex differences in neuroendocrine mechanisms mean that not only are behaviors often sexually dimorphic, but even if the sexes exhibit similar behaviors, a baseline difference in brain function may require distinct behavioral mechanisms (De Vries, 2004). In the monogamous oldfield mouse, *Peromyscus polionotus*, for example, genome-wide association studies implicate different mechanisms in the regulation of male and female parental care (Bendesky et al., 2017). The “dual function hypothesis” has suggested that similarly dimorphic mechanisms may underlie bonding or parental behaviors more generally (De Vries, 2004). To understand attachment, it is essential to examine neural function across the entire brain (López-Gutiérrez et al., 2021; Yee et al., 2016), and to explore individual and sex differences in such circuits as bonds form.

To systematically examine the mechanisms of pair-bond formation, we developed a whole-brain imaging and computational analysis pipeline that includes the first 3D histological atlas of the prairie-vole brain. This atlas and analysis pipeline enables the high-throughput, automated counting of cell markers throughout the prairie-vole brain. We used this tool to test for sexual dimorphism in the structure of prairie-vole brains, and to compare the gross anatomy of the prairie vole brain to the laboratory mouse. In order to map regions and circuits active during bonding, we next quantified immediate-early gene (IEG) induction in 824 brain regions at key times in bond formation. This represents the first unbiased identification of circuits active during bonding. Doing so allowed us to examine how the experience of mating finds its way into bonding circuits. It also allowed us to rigorously test two alternative hypotheses: that mating and bonding promote coordinated changes in the brains of mated pairs; and conversely, that sexually dimorphic circuits underlie bonding in males and females.

## Results

### A novel whole-brain imaging pipeline for the prairie vole

We began by generating a common coordinate framework (CCF) for the prairie-vole brain by iteratively averaging tissue autofluorescence from 191 brains imaged using light-sheet fluorescence microscopy (LSFM) (Video 1). Each of the brains were co-registered into this coordinate framework for computational analyses. Next, we registered an LSFM-based CCF of the mouse brain onto the prairie-vole CCF, enabling us to apply anatomical labels of the Allen Reference Atlas (ARA; Figure 1A, Figure 1—figure supplement 1, and Video 1) (Dong, 2008). This alignment revealed that the prairie-vole brain is ∼30% larger than the mouse brain and distinct in shape, but that relative volumes of brain regions were consistent across species and showed no evidence of sexual dimorphism (Figure 1A, Figure 1—figure supplement 1, Supplementary File 1, and Supplementary File 2).

**Figure 1.**
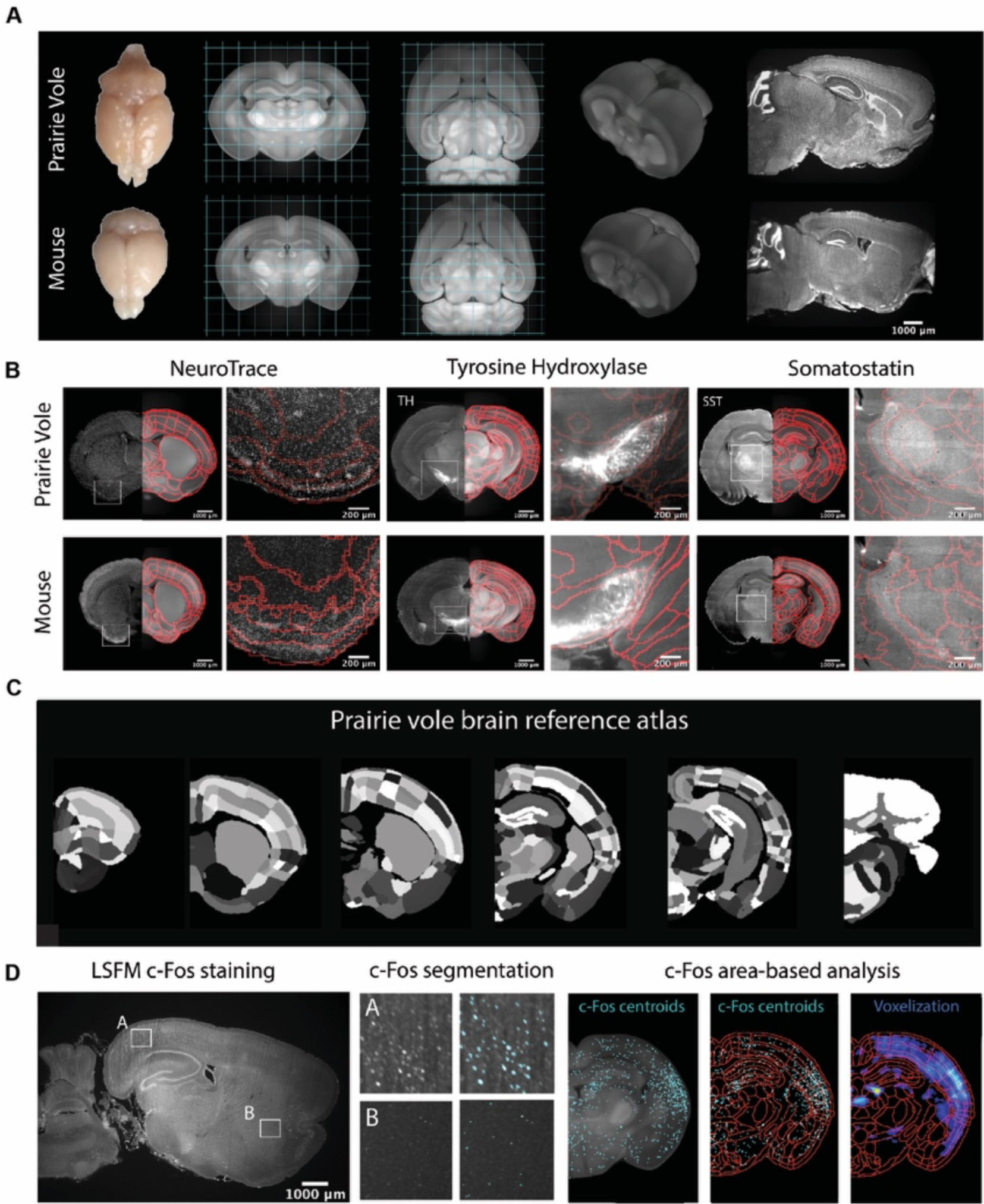
Prairie vole reference brain and atlas generation for automatic c-Fos analysis. **(A)** Generation of prairie vole (upper row) and mouse (lower row) reference brains using LSFM imaging. Shown is the top view of both prairie vole and mouse brains after perfusion. Also shown are cross-sections of coronal and horizontal views of prairie vole and mouse reference brains built after the co-registration of ∼200 brains per species. The prairie vole whole brain was 1.39 times bigger than the mouse (see Figure1—figure supplement 1 and Supplementary File 1). Both prairie vole and mouse brains underwent 3D renderization. Also shown are sagittal views of prairie vole and mouse whole brain fluorescent Nissl (NeuroTrace) staining imagined with LSFM. **(B)** Whole brain staining of both prairie vole (upper row) and mouse (lower row) brains registered to the reference brain. In red, boundaries of registered mouse reference atlas are plotted onto both prairie vole and mouse reference brains. NeuroTrace, TH and SST were registered and overlaid onto the atlases for validation. **(C)** Coronal sections of the resulting prairie vole atlas after manual validation. **(D)** Overview of LSFM prairie vole c-Fos+ analysis pipeline. Shown in the left panel is a sagittal section of prairie vole c-Fos immunolabeling imaged with LSFM. Shown in the center panel are detailed views of two brain locations immunolabeled with c-Fos and overlaid with resulting segmentation. Shown in the right panel is an area-based analysis of c-Fos+ cells. All c-Fos+ cells centroids are registered to the prairie vole reference brain and analyzed using the new prairie vole reference atlas. For each brain, a voxel representation is generated of all c-Fos+ cells in the same prairie vole reference space and overlay with the reference atlas.

To validate and refine the ARA anatomical borders in the prairie-vole brain, we performed whole-brain Nissl staining as well as iDISCO immuno-labelling targeting the cell-specific markers tyrosine hydroxylase (TH), parvalbumin (PV) and somatostatin (SST); we then aligned these images onto the prairie-vole CCF (Figure 1B,C and Figure 1—figure supplement 1). Lastly, we adapted for the prairie vole our computational pipeline for whole-brain detection and statistical comparisons of c-Fos+ neurons in LSFM-imaged brains (Kim et al., 2015, 2016; Renier et al., 2016) (Figure 1C). With a robust atlas and analysis tools in place, we conducted a detailed study of the time-course of mating and bond formation to examine IEG induction across the prairie-vole brain and test our hypotheses.

### Pair bonding involves a dynamic repertoire of social interactions

Sexually responsive voles typically mate within the first hour of pairing, and repeated mating over ∼6h initiates partner preferences characteristic of attachment (DeVries & Carter, 1999). The bond stabilizes between ∼12-24h, leading to prolonged changes in attachment and related behaviors (DeVries & Carter, 1999; Williams et al., 1992). With these milestones in mind, we precisely manipulated mating experience and examined how repeated sexual behaviors lead to a bond (Figure 2A, Figure 2—figure supplement 1, Figure 2—figure supplement 2).

**Figure 2.**
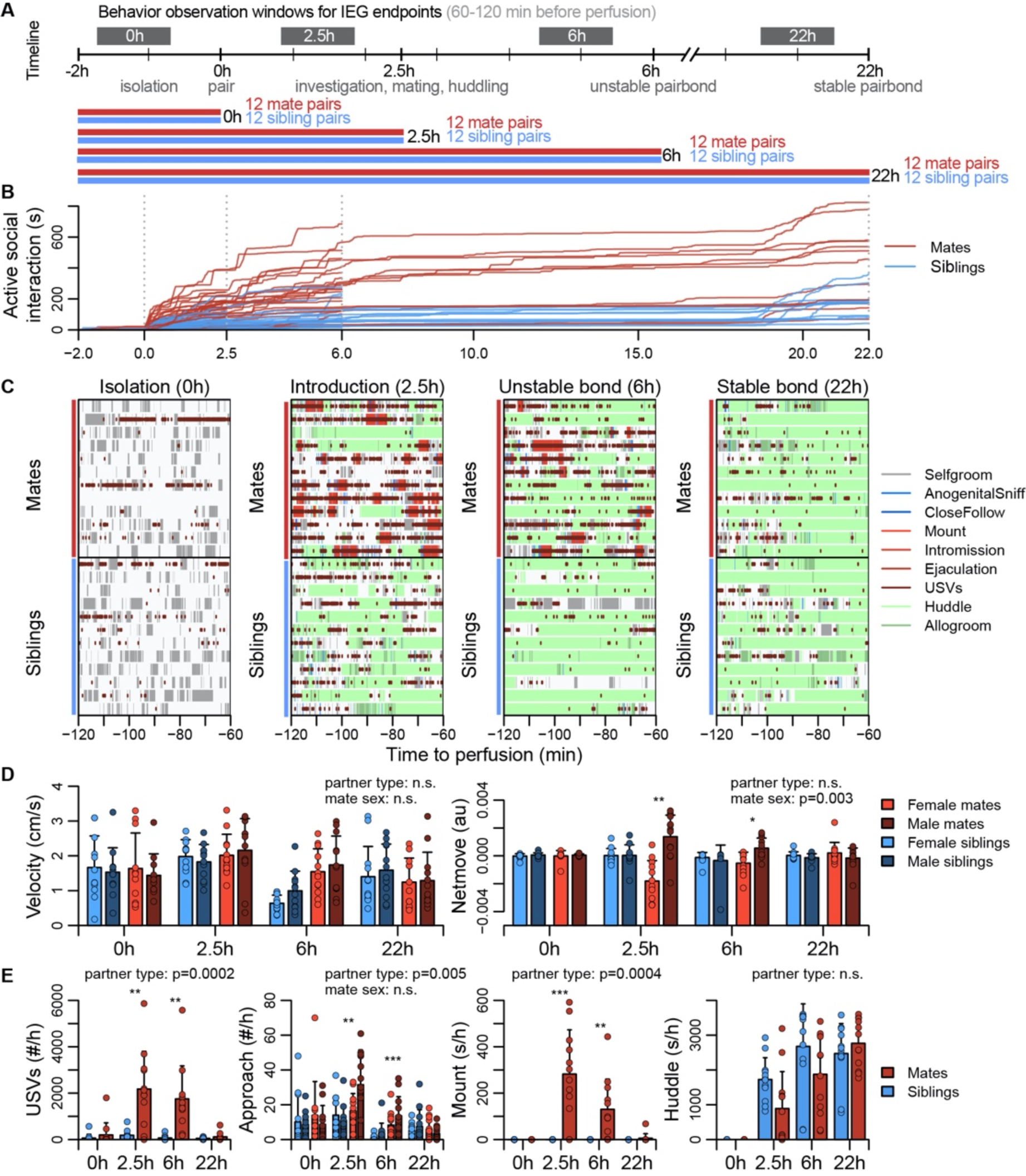
Study design and development of the prairie vole pair bond. **(A)** Schematic of the experiment design, where social behaviors (during 1h observations, black blocks) and IEG expression patterns are compared between mate pairs and siblings across time. **(B)** Continuous automated tracking of active social interactions (i.e., an index of social investigation and mating) for mate pairs (red lines) and sibling pairs (blue lines). **(C)** Time courses of specific social behaviors during 1h observation windows for IEG expression, with each row representing one pair. Social behaviors are overlaid onto self-grooming, and ultrasonic vocalizations (USVs, short red ticks) are overlaid onto all behaviors. **(D)** Plots showing group differences (mean ± sd) in individual activity level (velocity) and movement relative to the partner (net move, positive values indicate movement towards the partner). **(E)** Group differences (mean ± sd) in vocal behavior, proximity seeking, mating, and side-by-side contact. For (D) and (E) mate pairs are in red and sibling pairs in blue. Females are a lighter hue and males a darker hue for behaviors measured on an individual (rather than dyadic) level. T-tests were used to compare mates and siblings, and paired t-tests were used to compared female and male mates.

To coordinate the timing of mating, all subjects were isolated for 4-5 days, females were brought into estrus, and both sexes were screened for sexual receptivity (see Figure 2A). Subjects assigned to the bonding condition were paired with a novel opposite-sexed individual; to control for non-sexual social affiliation, remaining subjects were re-paired with a same-sex sibling who had been a cagemate prior to isolation. Members of each pair were isolated on either side of a divider for 2h. Following this acclimation, the divider was removed, and the pair could interact freely. (Figure 2A). Behavior sessions were terminated following a fixed timeline: animals were euthanized just before barrier removal (0h), following initial mating (2.5h), after initial bond formation (6h), or after bond stabilization (22h). Automated behavioral measures, such as proximity, vocalization, and relative movement were scored throughout the behavioral sessions. We performed detailed manual scoring for 1h focal intervals beginning 2h before euthanasia – a timeframe that reveals the behavioral states of animals during IEG induction (Figure 2B,C, Figure 2—figure supplement 1, Figure 2—figure supplement 2, Figure 2—figure supplement 3, and Supplementary File 3).

Mate and sibling pairs did not differ in locomotor activity or time spent in side-by-side contact (“huddling”; two sample t-tests: velocity, n = 94 mates and 96 siblings, t = 1.592, FDR-corrected q=0.170, huddling, n = 47 mate and 48 sibling dyads, t = 1.122 and 0.3663; Figure 2D). Mate pairs progressed through known stages of mating behavior, showing elevated rates of anogenital investigation and vocalization (t-tests: investigation, n = 94 mates and 96 siblings, t = 4.184, q=0.0003, vocalizations, n = 48 mate and 48 sibling dyads, t = 4.708, q=0.0003); males moved more often toward females, while females moved away from their partners (paired t-test: net movement, n = 47 females and 47 males, t = 3.127, q=0.0138), a behavior consistent with female copulatory pacing (Pfaus et al., 2001). Consummatory aspects of mating – mounting, intromission and ejaculation – were common in both the 2.5h and 6h focal windows; by 22h mating was rare, and levels of male-female huddling resembled those of same-sex siblings housed together since birth (Figure 2E). This profile is consistent with literature on the timing and process of prairie vole bond formation (DeVries & Carter, 1999; Williams et al., 1992).

### A brain-wide functional neural network for pair bonding reveals 7 major neuroanatomical clusters, with special prominence for regions of the BST and hypothalamus

Next we measured brain-wide c-Fos immunostaining, a common proxy for neuronal activity and plasticity (Sheng & Greenberg, 1990). To identify brain areas that differed between mate-paired and sibling controls, we used a generalized linear model (GLM) to compare two alternative models. A null model included terms for sex, time, and block; a full model included these variables, as well as pairing status and two interaction terms (pairing*sex, pairing*time). Brain-wide comparisons using voxels or regions of interest (ROI) were largely concordant (Figure 3A,B, Figure 3—figure supplement 1, and Video 2), revealing an extensive but specific network of brain regions active during mating and bonding. The brain atlas is organized hierarchically, and of 99 ROIs that differed significantly (permutation test: n = 189 animals, permutations = 10,000, FDR-corrected α < 0.1; Figure 3B and Supplementary File 4), 68 regions were anatomically distinct.

**Figure 3.**
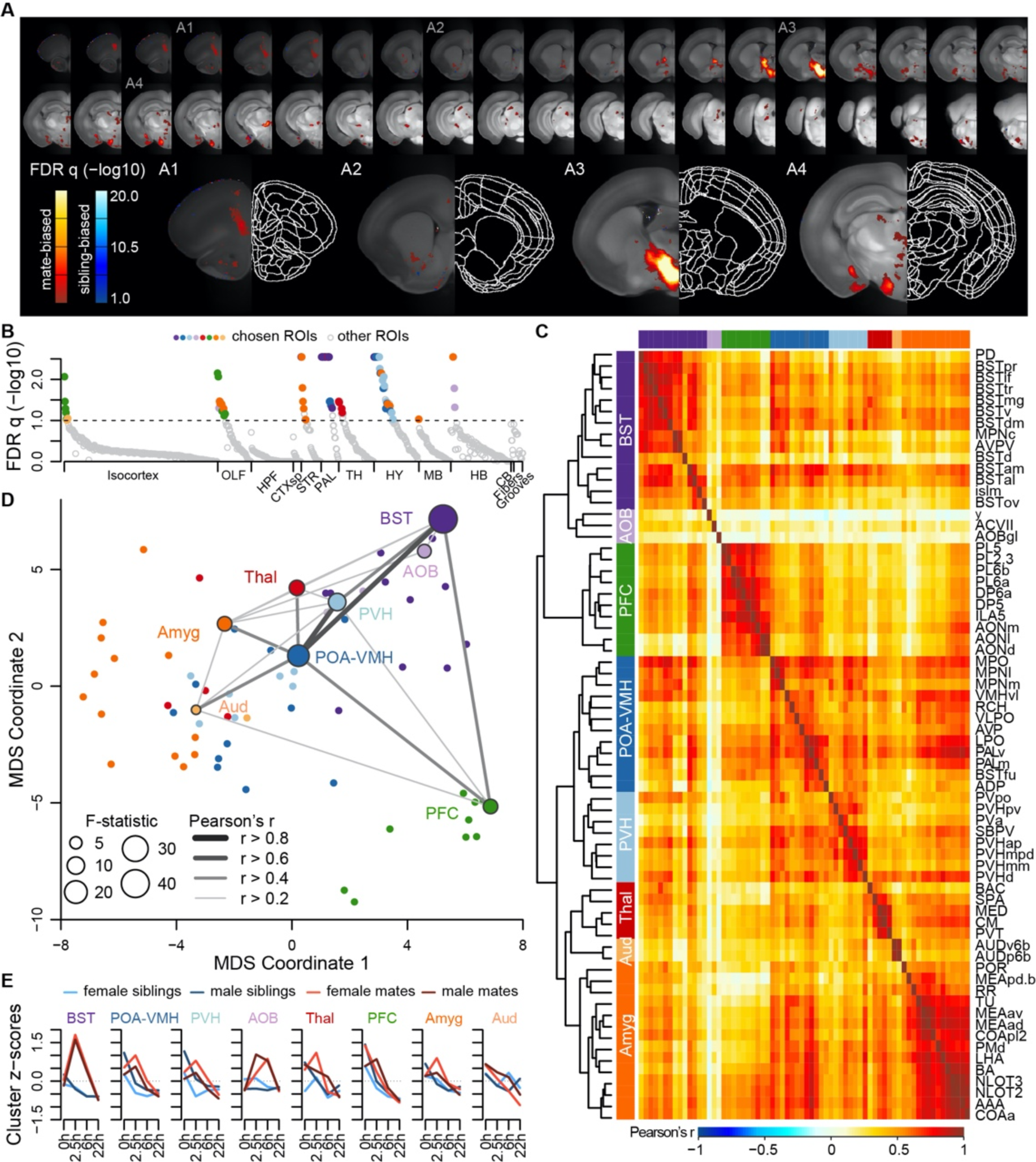
A brain-wide functional network is active during pair bond formation. **(A)** Map of bonding-associated voxels. Brightness indicates significance level of comparisons of hypothesized and null models (GLMs) to predict c-Fos cell counts. **(B)** Identification of significant and mutually exclusive ROIs, sorted by anatomical division and F-statistic. The significance levels of these models were determined with a permutation test, by comparing the observed F-statistic to a null distribution of F-statistics from shuffled data (n=10,000 permutations). Colored symbols for ROIs match their cluster group assignments in 3C and 3D. **(C)** Hierarchical clustering of chosen ROIs (n=68) and pairwise Pearson correlations of c-Fos cell counts. **(D)** Multi-dimensional scaling (MDS) coordinate space of the correlations between chosen ROIs. The most significant ROIs per cluster are labelled and their symbol size scaled by the F-statistic. Darkness and thickness of connecting lines reflect correlation coefficients. **(E)** Time course trajectories of total c-Fos cell counts within each cluster. Counts per cluster are scaled across samples and averages taken for each experiment group, with red lines for mates and blue for siblings (females-lighter, males-darker). Each cluster is given labels to summarize the most significant ROIs within them.

A hierarchical cluster analysis assigned these 68 ROIs into 8 groups (Figure 3C,D, Figure 3—figure supplement 2, and Figure 3—figure supplement 3). Each of these groups, or ‘clusters,’ included multiple brain regions but often centered on a specific structure or group of structures. The purple “BST” cluster, for example, contained regions of the preoptic area and periventricular nucleus of the hypothalamus, as well as multiple substructures within the bed nucleus of the stria terminalis. The blue “POA-VMH” and light blue “PVH” clusters contained multiple regions within the hypothalamus, including the paraventricular nucleus of the hypothalamus, as well as regions of the preoptic area. The green “PFC” cluster was composed of regions within both the prefrontal cortex and olfactory areas. The orange “Amyg” cluster contained the lateral hypothalamus and various olfactory related regions, but the majority of the cluster consisted of amygdalar nuclei such as cortical and medial amygdala and the anterior amygdala area. The light orange “AUD” cluster contained some regions within auditory cortex, and the red “Thal” cluster involved a variety of thalamic regions. The light purple “AOB” cluster contained minimally correlated regions such as the accessory olfactory bulb, and so we do not consider this group to be one of the major clusters. Comparing the groups with connectivity reported in the ARA mouse ‘connectome’ (Knox et al., 2019) suggests that 6 major clusters in our dataset (BST, POA-VMH, PVH, PFC, Amyg, and AUD) are anatomically connected (permutation test: permutations = 10,000, p=0.0001, Figure 3—figure supplement 4).

Although the nucleus accumbens did not survive multiple test corrections in our ROI analysis (n = 189 animals, F = 2.936, q=0.1747), it was significant in the univariate analysis (p=0.0304), particularly when focused on the 2.5 and 6h timepoints (two sample t-test: n = 47 mates and 47 siblings, t=2.530, p=0.0138, Video 2). Furthermore, voxel-level comparisons revealed significant sites within the ventral striatum and the posterior nucleus accumbens (Figure 2A, Figure 3—figure supplement 1, and Video 2).

### Major dimensions of neural and behavioral variation are coordinated among mated pairs

Canonical correlation analysis (CCA) reveals the latent correlational structure within two sets of variables, and so is well suited to compare the principal dimensions of behavioral variation to its neural counterparts (Figure 4, Figure 4—figure supplement 1, and Figure 4—figure supplement 2). We found that the first canonical correlate (CC1), which defines the largest axis of shared variation in brain and behavior, loaded highly on the BST, POA-VMH, and PVH clusters, as well as on mating-related behaviors (Figure 4A-D). CC1 scores captured responses to mating and bonding in the 2.5h and 6h timepoints (Figure 4A,B). The second canonical correlate (CC2) captured differences between animals who were isolated or paired and loaded particularly highly on the limbic cortical cluster (PFC; Figure 4B, Figure 4—figure supplement 1).

**Figure 4.**
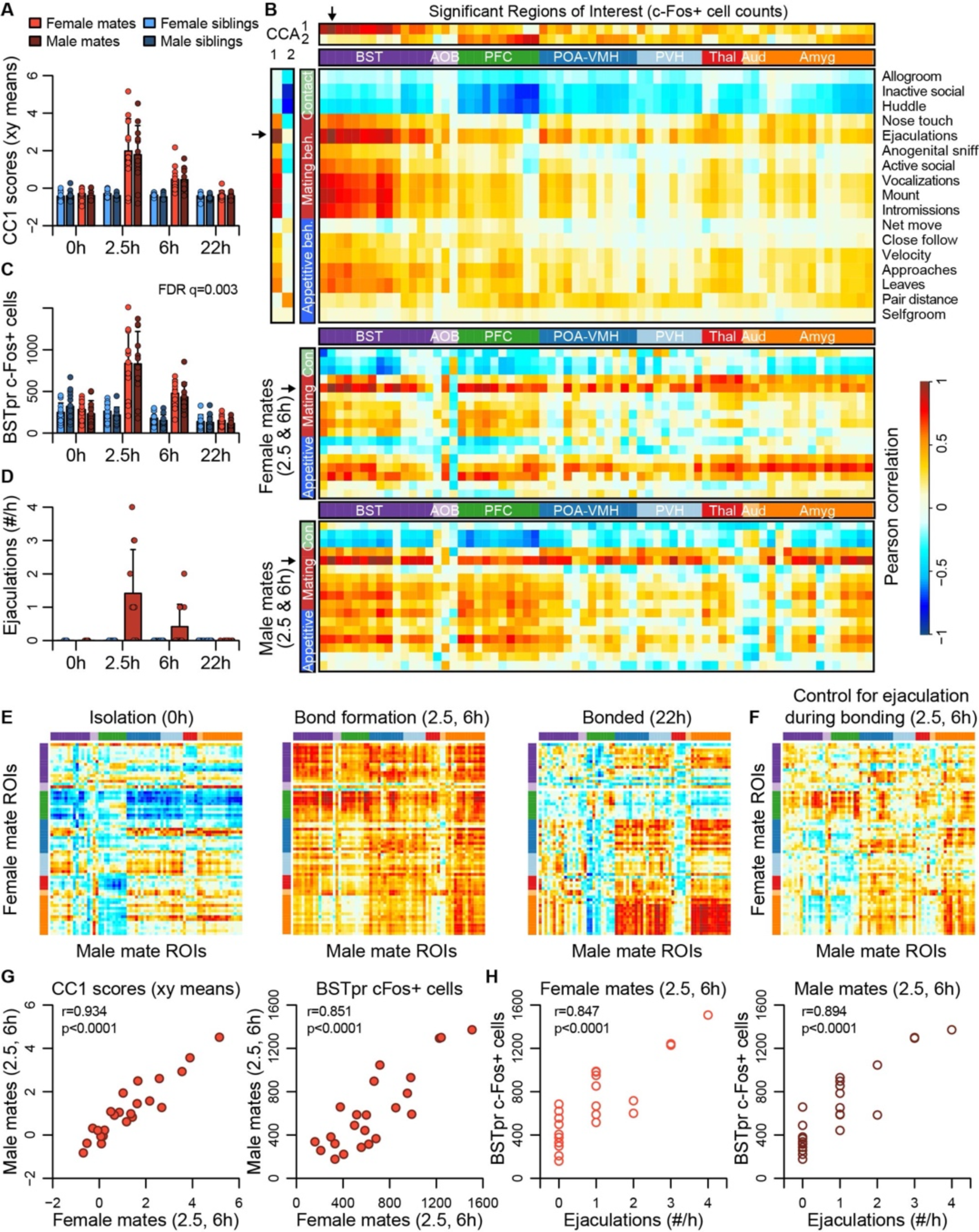
BST emerges as a central hub in the bonding network that is associated with mating success in both sexes. **(A)** The first dimension of canonical correlation (CC) scores is compared across experiment group (mean ± sd). **(B)** Heatmaps represent correlation coefficients among CC scores, ROI c-Fos cell counts, and behavior measures. The full dataset is on top, and the bottom two correlograms are for female and male mates. For (B), (E) and (F), warm and cool colors represent positive and negative coefficients, respectively. Arrows mark the ROI (BSTpr) and behavior (ejaculation rate) with the strongest correlations to CC1. **(C)** BSTpr activation is compared across groups with a model comparison followed by a permutation test. **(D)** Successful mating events are shown across timepoints (mean ± sd). **(E)** Similarity (i.e., correlation) of bonding network activation is shown in female-male pairs. **(F)** Similarity in bonding network activation is shown in female-male pairs, using partial Pearson correlations to control for ejaculation rate. **(G)** Female-male pair similarity is shown for CC1 scores and BSTpr activation during bond formation. **(H)** Mating success is associated with BSTpr activation for both sexes during bond formation. Pearson correlations were used to assess these associations.

The first two CC dimensions showed no evidence of sexual dimorphism, while the third dimension suggested subtle but non-significant differences between male and female mates (Figure 4—figure supplement 1). A formal two-model comparison isolating the effect of the sex*pairing interaction revealed no ROIs that differed following FDR correction. If we limit the comparison to just those 68 areas identified as responding to pairing, we find 29 unique ROIs that exhibit evidence of sex*pairing interaction, although these effects are much weaker than those identified in our above analysis of pairing (Video 2).

Although differences between males and females were modest, the CCA suggested substantial individual differences among mated pairs. We used these data to test whether pairing involved a coordinated change in activities across brain regions. We find that during mating and bond formation (2.5-6h), activity is strongly correlated across ROIs (Figure 4E,F and Figure 4—figure supplement 3). Similar to these patterns distributed across putative pair-bonding regions, CC1 brain scores show a strong correlation between male and female mates (Pearson correlation: df = 22, r=0.93, p<0.0001, Figure 4G).

Following bonding (22h), within-pair correlations are confined to the coupling of activity between the hypothalamus and amygdala, including between MPOA and medial amygdala. In siblings, who were housed together since birth, hypothalamus-amygdala correlations are relatively uniform across time points, while correlations outside of these regions are largely absent (Figure 4—figure supplement 3).

The principal nucleus of the BST (prBST), a subregion of the posterior BST, loads highly on CC1 (Figure 4B), and exhibits a strong response to mating in the 2.5 and 6h time points (permutation test: n = 189 animals, permutations = 10,000, q=0.003, Figure 4C); we found that this brain region also shows strong correlations in heterosexual pairs (Pearson correlation: df = 21, r=0.851, p<0.0001, Figure 4G). Remarkably, the number of observed male ejaculations is strongly predictive of prBST c-Fos+ cells in both males and females (males: df = 22, r=0.894, p<0.0001; females: df = 21, r=0.847, p<0.0001; Figure 4E). To our surprise, but consistent with the canonical correlation analysis, male ejaculation rates were the strongest predictor of activity across brain regions of both sexes (Figure 4D). Statistically controlling for the number of ejaculations effectively abolished the correlation we observed in mated pairs (Figure 4F).

## Discussion

Here, we present a brain-wide network of IEG+ circuits active as mating experience elicits a pair bond. We find no evidence of anatomical sexual dimorphism, and only modest evidence of dimorphic function. Our whole-brain mapping of pair-bond formation implicates 68 unique brain regions, including 18 regions that are within the primary brain network proposed for prairie-vole bond formation (Walum & Young, 2018). The 68 identified regions are more strongly anatomically connected to one another than predicted by chance, and so can be interpreted as circuits active during pairing.

Although the majority of regions identified have not been linked directly to bonding, many are logical components of a pair-bonding network. For example, the strongest effects of pairing were detected within a cluster containing multiple compartments of the posterior bed nucleus of the stria terminalis (BST), as well as the posterodorsal preoptic nucleus (PD), core medial preoptic nucleus (MPNc) and anteroventral periventricular nucleus (AVPV; “BST” purple cluster, Figure 3C-E and Figure 4A-C). While this cluster exhibited the strongest response to pairing, only the BST has been previously implicated in prairie-vole bonding (Cushing & Wynne-Edwards, 2006; Lei et al., 2010). The cluster as a whole, however, precisely matches the neural circuitry of male ejaculation previously mapped in rats, gerbils and hamsters (Coolen, 2005; Heeb & Yahr, 1996, 2001; Pfaus, 2009; Simmons & Yahr, 2002). The posterior BST projects to both the medial preoptic nucleus and the paraventricular nucleus, major contributors to two other clusters (“POA-VMH,” blue; “PVH,” light blue), regions that also exhibit strong responses to pairing (Cushing et al., 2003; Insel & Young, 2001). The paraventricular nucleus of the hypothalamus, moreover, is a major source of the neuropeptides oxytocin and vasopressin, known modulators of pair bonding in the prairie vole (Walum & Young, 2018).

A fourth cluster (“PFC,” green) is composed of prelimbic, infralimbic and olfactory cortex; activity in the vole prefrontal cortex is known to be modulated by hypothalamic oxytocin, and to shape bonding through projections to the nucleus accumbens (Amadei et al., 2017; Burkett et al., 2016; Horie et al., 2020). The pattern of activity in this cluster, however, indicates that it was due in part to differences between the isolated animals (0h) and other time points (Figure 4—figure supplement 1 and Figure 4—figure supplement 2). Because animals in the isolated condition were in a compartment adjacent to either an opposite sexed individual or a familiar former cagemate, we cannot rule out that olfactory or auditory cues may have made animals aware of the presence of a potential social partner. Indeed, we interpret this dimension as capturing appetitive aspects of behaviors associated with investigation of the animal isolated from the subject by the barrier.

Although the PFC and other olfactory cortical areas formed a cluster, we did not find widespread c-Fos induction throughout the cortex in response to pairing. It seems likely that sensory and motor areas were important for social processes related to both pair-bonding and reunion with same-sex cagemates, such as investigation and recognition. Our study design, however, highlights differences between treatments, and in order to detect such effects, it might be necessary to compare mating and bonding pairs to animals left in complete isolation. Moreover, several cortical regions that did not survive corrections for multiple tests may have been identified in a less stringent analysis. Several subregions within the isocortex, hippocampal formation, and cortical subplate had statistical models that approached significance (i.e., p-values < 0.1) prior to multiple test corrections. These subregions were found within primary somatosensory area, primary auditory area, dorsal and ventral auditory areas, primary visual area, anteromedial visual area, agranular insular area, temporal association areas, ectorhinal area, postsubiculum, and basomedial amygdala. Frontal cortex subregions were within the agranular insular area and orbital area, as well as additional subregions in prelimbic and infralimbic areas of the PFC.

In addition to its effects in the PFC, pairing drove increased c-Fos expression in the ventral pallidum, a major node in reward circuity, as well as in the paraventricular nucleus and the medial preoptic area, modulators of reward. This is consistent with a large body of work implicating neuropeptide actions on reward circuits in the formation of bonds (Walum & Young, 2018; Young & Wang, 2004). Conspicuously missing from our list, however, is significant pairing-induced c-Fos induction in the nucleus accumbens. The absence of significant accumbens IEG induction may reflect the limitations of using c-Fos and other immediate early genes as indicators of neural activity. It is known that some neuronal populations can be active without expressing c-Fos (Sheng & Greenberg, 1990). Indeed, although a variety of studies implicate the accumbens in bond formation (Amadei et al., 2017; Aragona et al., 2006; Scribner et al., 2020), previous work finds only weak c-Fos induction in the prairie vole accumbens during bonding (Curtis & Wang, 2003). Another possibility is that there was heterogeneous activation in the accumbens that was not captured by the precision of our atlas. Consistent with this interpretation, found that the accumbens was significant in univariate tests, as well as in voxel-level analyses. Overall, our results do not conflict with pharmacological, electrophysiological, and calcium-imaging data on the role of the nucleus accumbens in prairie vole bonding (Amadei et al., 2017; Aragona et al., 2006; Scribner et al., 2020). Instead, the absence of significant effects at the level of the entire nucleus accumbens together with the presence of anatomically restricted voxel-level significance suggests substantial anatomical heterogeneity in the contributions of the nucleus accumbens to bond formation.

Our data suggest that pairing-related c-Fos immunoreactivity is largely shared across sexes, and that much of the pairing related activity is driven by mating behavior. The strongest signal of pairing status was in the prBST, a region important in male ejaculation (Insel & Young, 2001; Pfaus, 2009). We found that both sexes exhibited the strongest signal of pairing status in prBST. This finding supports recent work in mice showing that different neural populations in prBST, aromatase+ and Vgat+ neurons, respond to ejaculation in males and females (Bayless et al., 2019). Moreover, we found a surprisingly strong relationship between ejaculation rates and brain-wide activity patterns across the putative pair-bonding circuit. The concordance between males and females was surprising, but is consistent with IEG studies of ejaculation and related circuitry in male and female rats (Coolen, 2005; Pfaus, 2009). These data are also consistent with the interpretation that copulation enables coordination and assessment during bonding (Carter et al., 1995; Dewsbury, 1988; Eberhard, 1996; Getz et al., 1993). Our findings, along with this previous work, support the hypothesis that sexual behavior plays a key role in driving pair-bond strength. However, the current study focused on animals that were screened for sexual receptivity, which may have limited variation in sexual behavior across pairs. An intriguing direction for future research will be to test how this variation contributes to bond strength.

The tight coupling of widespread neural activity adds to the growing number of examples – including fish, mice, bats, and humans – that demonstrate correlated neural function across socially interacting individuals (Hasson et al., 2012; Kingsbury et al., 2019; Kinreich et al., 2017; Long et al., 2020; Vu et al., 2020; Zhang & Yartsev, 2019). Although widespread differences among mated pairs seem to be driven by ejaculation rates during mating and bonding, a smaller subset of circuits – specifically connections between the hypothalamus and amygdala – remain correlated after pairing. The similarity of this post-pairing pattern to non-sexual affiliative mechanisms recently documented in lab mice (Hu et al., 2021), and to correlations we observe among sibling pairs, suggests these regions play a role in both reproductive and non-reproductive attachment. These brain regions, and especially the amygdala, will be important candidates for future research on neural regulation of pair-bond maintenance.

Before offering a synthesis of our findings, it would be useful to acknowledge or reiterate a few caveats. First, as noted above, IEG induction does not capture all relevant neural activity (Sheng & Greenberg, 1990). Second, the design of our experiment, which controlled for social interaction, likely excluded many circuits important to both pair bonding and sibling social interactions. Third, c-Fos activity within a given brain region may nevertheless rely on distinct cell types, and so the absence of sex differences in c-Fos immunoreactivity does not definitively rule out the sexually dimorphic circuits hypothesized in the “dual function hypothesis” (De Vries, 2004). Lastly, the current study focused on animals that were screened for sexual receptivity, which may have limited the variation in sexual behavior across opposite-sex pairs.

### Ideas and speculation

This brain-wide map of IEG induction provides a uniquely comprehensive perspective on the circuits that enable sexual behavior to become a bond. To help organize this enormous dataset, we looked to existing literature to compare the implicated circuits to those related to sociosexual behaviors in other species (Figure 5A,B). Overlaying the time course of behaviors and circuit activity suggest distinct stages of bonding and attendant neural function. We divide those stages into mating, bonding, and ongoing bond maintenance.

**Figure 5.**
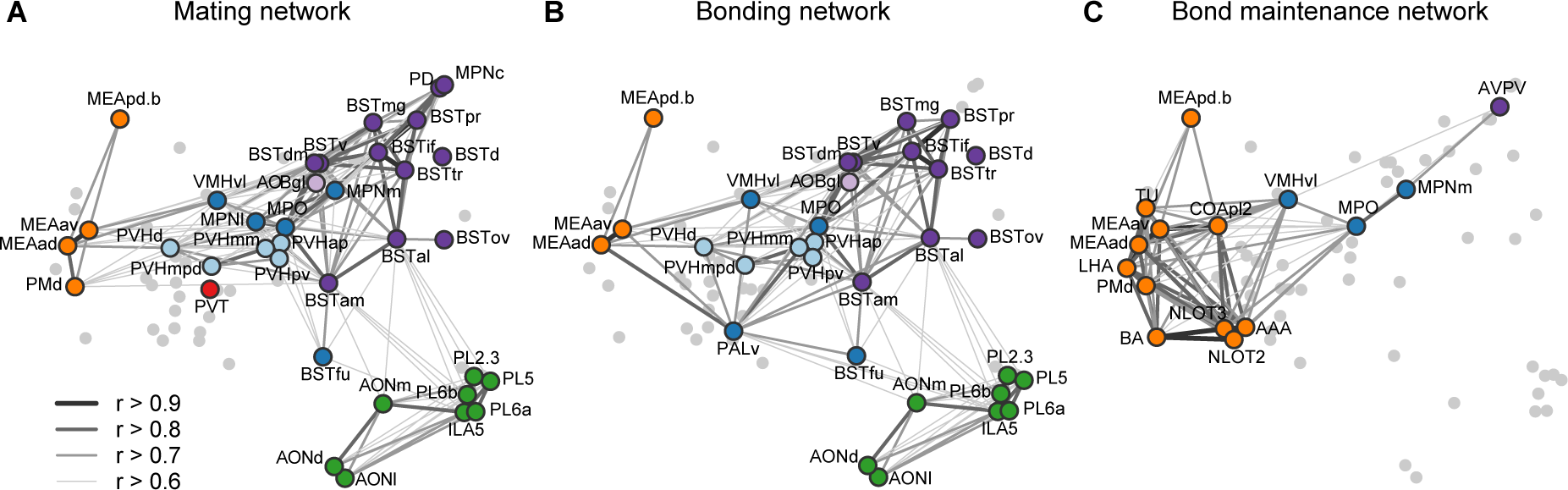
Working model for neural systems that shape stages of pair bond development. **(A)** Schematic of the network of regions identified in our study overlaid with regions known to be involved in rodent mating behavior (Pfaus & Heeb, 1997; Veening et al., 2014; Veening & Coolen, 2014). This network is proposed to be involved during the early stages of bond formation (e.g., 2.5 h timepoint). **(B)** Schematic of the network of regions identified in our study overlaid with regions known to be involved in prairie vole pair bonding behavior (Walum & Young, 2018; K. A. Young et al., 2011). This network is proposed to be involved with the middle stages of bond formation when animals are engaged in prolonged mating and affiliative interactions (e.g., 6 h timepoint). **(C)** Schematic of the network of regions that are correlated between female and male mates at the 22 h timepoint in this study. These are identified from pairs of regions in which both sexes show high inter-individual similarity (Pearson correlation, r > 0.75). This network is proposed to be involved with the recognition and convergence of behavioral state in bonding partners. The network schematics in (A), (B) and (C) are adapted from the multi-dimensional scaling of correlations between region activity (Figure 3D). Light gray points represent regions not included in the proposed network. Line thickness and darkness between colored regions represent Pearson correlation r values between the connected regions.

The first canonical correlate captured the major dimensions of both brain and behavioral variation, and showed the largest group differences surrounding the 2.5 hr timepoint. Behaviorally, this difference mapped strongly onto mating behavior, and in particular to rates of male ejaculation. The neural circuits of rodent sexual behavior in general, and of male ejaculation in particular, have been well studied in rats, hamsters and laboratory mice (Bayless et al., 2019; Pfaus & Heeb, 1997; Veening et al., 2014; Veening & Coolen, 2014). In this context, chemosensory signals are thought to drive activity in MeA, MPO, and PVH. Copulation drives activity in MPN, PD, VMH and BST. We found that elevated c-Fos induction levels occur in these areas early on in the bonding timecourse and then decline. Such findings support the idea that these core circuits orchestrate early social interactions that enable individuals to process chemosensory signals and then initiate lordosis, mounting, and ejaculation. Our study also revealed a set of regions that have yet to be emphasized in socio-sexual circuits, including lateral hypothalamic area and lateral preoptic area. While research on these regions emphasize non-social functions, a number of studies suggest that these regions play a role in social behaviors, or participate in goal-directed behavior and reward (J. Chen et al., 2021; Nieh et al., 2016; Petzold et al., 2023).

Interestingly, one anomaly from the literature on sexual behavior circuits is that females also show IEG induction associated with male ejaculation – a finding evident in both rats and mice (Pfaus & Heeb, 1997; Veening et al., 2014). Similarly, recent work in female laboratory mice demonstrate that Vgat+ neurons in the prBST exhibit responses to male ejaculation (Bayless et al., 2019). The extraordinary pattern of coordination we observe between members of a pair across nearly 70 brain regions, and its prediction by male ejaculation, suggest that both males and females are experiencing similar and profound affective states. Moreover, the fact that repeated copulation is necessary for bonding is consistent with work in a variety of taxa suggesting that repeated copulation is a means of partner assessment (Dewsbury, 1988; Eberhard, 1996); in prairie voles, we suggest that both males and females are assessing the ability of a male to monopolize a female, a trait that would predict male paternity and ability to defend a nest against conspecifics in the field. Pfaus and others have argued that females of other species, including laboratory rodents, exhibit orgasm-like responses (Georgiadis et al., 2012). Although our current data are unable to address this claim directly, the hypothesis offers a parsimonious interpretation of our data, and the topic merits further scrutiny.

In this circuitry, the “extended amygdala” regions of the BST and MeA stand out for their extensive projections to and from the hypothalamus, and for their known roles in individual recognition. Dumais et al (2016), for example, have documented that the BST of the rat is essential to individual recognition; in parallel, Knoedler et al (2022) find that ERa in the BST coordinates sex recognition in laboratory mice. Most directly, Cushing and colleagues have shown that ERa in the BST is essential to the formation of pairbonds in prairie voles (Lei et al., 2010). Similarly, oxytocin actions in the MeA are known to govern social recognition in lab mice (Ferguson et al., 2001). We hypothesize that IEG induction in response to mating enables the identity of a mate to get access to the modulation of hormonal and behavioral states by the hypothalamus. Such identity representations may coordinate partner-specific patterns of neuropeptide release (PVN), selective aggression (VMHvl), selective mating (VMHvl, MPOA), and social reward (MPOA).

After bond formation and stabilization, we find that that mated pairs are similar to co-housed siblings in terms of brain-wide IEG induction patterns (Figure 5C). More remarkable, is that after 22 hrs both co-housed siblings and mated pairs show strong correlations in amygdala IEG induction, particularly in the MeApd. We propose that this correlated activity reflects a sensitivity to the behavioral state of a partner that emerges as a function of bonding. Recent work has revealed that in laboratory mice, tachykinin+ GABAergic neurons project from the MeApd to the MPOA, where they contribute to both consolation grooming and social reward (Hu et al., 2021; Wu et al., 2021). These projections have not been implicated in pairbonding, but our current data suggest this circuit may play a broader role in the ongoing function of the relationships of both mates and siblings. Since prairie voles often live in larger family groups (Getz et al., 1993), particularly over winter, a common mechanism for the maintenance of familial and sexual bonds would be consistent with their ecology. In addition, the strong correlation of activity across a variety of related amygdala and associated regions suggest this circuit may be anatomically much broader than the interactions between MeApd and MPOA documented in laboratory mice.

### Conclusions

Overall, our data survey the brain to identify circuits active as mating produces a pairbond. These data allowed strong tests of opposing hypotheses about sexual dimorphism and coordination during bonding (De Vries, 2004; Hasson et al., 2012; Kingsbury et al., 2019; Kinreich et al., 2017; Long et al., 2020; Zhang & Yartsev, 2019), revealing a surprising absence of sexual dimorphism in structure and function, and an extensive coordination of neural activity among new pairs. We confirm the activity of known regulators of mating and bonding such as reward circuits, the paraventricular nucleus and the bed nucleus of the stria terminalis (Coolen, 2005; Insel & Young, 2001; Lei et al., 2010; Pfaus, 2009); we map a path by which sexual activity finds its way to known regulators of bonding, and identify a variety of novel regions and circuits whose roles in bonding remain to be examined. Our results suggest a novel model in which the BST is a key node connecting sexual experience to the neuroendocrine functions of the hypothalamus and preoptic area, and that coupling between the preoptic area and the amygdala may play an unappreciated role in the maintenance of an established bond. Manipulations of these circuits and their behavioral consequences offer rich new opportunities for research into the mechanisms of bonding and their contributions to well-being.

## Materials and Methods

### Animals

Prairie voles (*Microtus ochrogaster*) used in behavioral and anatomical experiments were prairie voles derived from wild-caught voles from Jackson County, Illinois and bred at The University of Texas at Austin. At weaning (PND 21), voles were housed in polycarbonate cages (R20 Rat Cage, Ancare Corp., NY) in groups of 2-5 same-sex littermates and provided with standard rodent chow (LabDiet 5001, Lab Supply, TX) and water *ad libitum*. Temperature of the colony room was maintained at a controlled temperature (20-23°C), and the photoperiod was on a 12:12 light:dark cycle (lights on: 0600, lights off: 1800). Housing arrangements allowed animals to receive visual and olfactory cues, but not tactile contact, with conspecific males and females. Mice used for anatomical research were housed with ad libitum access to food and water in a controlled temperature (21-22°C) and light (12:12 light/dark cycle) room. All animal procedures were approved by the Institutional Animal Care and Use Committees at the University of Texas at Austin and Cold Spring Harbor Laboratory.

### Brain tissue sample preparation and processing

Prairie voles underwent intracardiac perfusion ∼20 min (mean = 21 min, range = 10-40 min) minutes after the behavioral observation endpoints (2h acclimation or 2h acclimation in addition to 2.5h, 6h or 22h cohabitation). Animals were anesthetized with open-drop isoflurane exposure, exsanguinated with 0.9% saline (2 min 30 s at 13 mL/min), and fixed with 4% paraformaldehyde (PFA) in 0.05M PB (5 min 30 s at 13 mL/min). All harvested brain tissue samples were post-fixed overnight at 4° C in 4% PFA in PB. After post-fixation, samples were washed 3x in 0.05M PB and stored in 0.05M PB 0.02% sodium azide at 4° C until immunolabeling processing.

Mice of 8-10 weeks old were anesthetized with ketamine/xylazine and transcardially perfused with isotonic saline followed by 4% PFA in 0.1M phosphate buffer (PB, pH 7.4). Brains were extracted and post-fixed overnight at 4° C in the same fixative solution, and stored at 0.05M PB until immunolabeling processing.

### Whole brain IEG staining and imaging

The right hemisphere of all brain samples were cut and immunolabeled for c-Fos and posteriorly cleared using iDisco+ protocol (Renier et al., 2014, 2016). Briefly, samples were initially delipidated with methanol and later permeabilized and blocked with DMSO and donkey serum respectively. Thereafter, all samples were incubated with c-Fos antibody and Alexa Fluorophore 647 secondary antibodies. Samples were cleared with increasing concentration steps of methanol and dichloromethane as previously described (Renier et al., 2014, 2016). After clearing, samples were imaged sagittally on a light-sheet fluorescence microscope (Ultramicroscope II, LaVision Biotec). Samples were imaged continuously every 5um at 640 nm and 488 nm, for signal and background channels respectively.

### Construction of the vole reference brain and atlas

For the construction of the prairie vole reference brain, 190 brains from c-Fos+ cell counting analysis were co-registered as described below (3D registration of the vole brain) (Kim et al., 2015). For the construction of the prairie vole reference atlas, the Allen Reference Atlas (ARA) was initially used as a template. After initial registration, output transformations were used to warp mouse atlas onto the prairie vole reference brain. In order to validate atlas registration, prairie vole Neurotrace stained brains were registered at high resolution onto the prairie vole reference brain.

### 3D registration of the vole brain

Brains were registered to a standardized reference brain as previously described (Renier et al., 2014, 2016). Initial 3D affine transformation was calculated using 6 resolution levels followed by a 3D B-spline transformation with 3 resolution levels. Similarity was computed using Advanced Mattes mutual Information metric by Elastix registration toolbox. In order to enhance image registration, both brain images and reference brains were pre-processed to reduce the impact of imaging artifacts during the computation of mutual information (Video 1). First, brain image illumination was corrected to homogenize illumination across sections. Second, both brain images and reference brain intensities were smoothed to reduce imaging artifacts (Muñoz-Castañeda & Osten; manuscript in preparation).

### Whole brain prairie vole Neurotrace staining

Whole brain Neurotrace staining was performed with a modification of the iDISCO+ protocol (Muñoz-Castañeda & Osten, manuscript in preparation) (Muñoz-Castañeda et al., 2021). Samples were initially washed in phosphate-buffered saline (PBS) and incubated for in PBS + TritonX-100 + DMSO + glycine. Then samples were transferred and incubated in a solution with PBS and Neurotrace. Finally, samples were washed in PBS.

### Comparison of vole and mouse neuroanatomy

For area-based volume quantifications, the reference brain was registered onto each single imaged brain and all volume areas were automatically quantified with custom made scripts (Muñoz-Castañeda & Osten, manuscript in preparation). These area volumes were used for comparisons within and between species.

### STPT whole brain imaging for anatomical delineation

Before imaging, both prairie vole and mouse brains were embedded and cross-linked with oxidized 4% agarose as previously described (Kim et al., 2017; Ragan et al., 2012). Whole brain imaging was achieved using the automated whole-mount microscopy STPT. The entire brain was coronally imaged at an X,Y resolution of 1µm and Z-spacing of 50µm (Kim et al., 2017; Ragan et al., 2012). After imaging, brains were registered to the reference brain for anatomical validation (see 3D registration of the vole brain).

### Automated c-Fos+ cell detection

c-Fos+ cells segmentation was performed using convolutional neural networks as previously described (Kim et al., 2016, 2017). After cell segmentation, all cell centroids were calculated for whole brain distribution analysis. For the analysis of area-based c-Fos+ cell counts, the mouse reference brain was registered onto each prairie vole brain, using the output transformations to warp the atlas onto each individual brain for ROI cell counting and distribution. For voxel analysis, the inverse transformation output was used to move all c-Fos+ centroids onto the reference brain.

### Experiment design

Study subjects were 8 to 12 week old sexually naïve prairie voles. There were eight treatments that varied based on partner type and cohabitation time (Figure 2). Subjects were partnered with a familiar same-sex cage mate (“siblings”) or an opposite-sex individual (“mates”) for 0, 2.5, 6, or 22 hours. The 0 h timepoint represents a baseline state before pairing takes place, the 2.5 h time point is ∼2 h after the first mating bout, the 6 h time point is when an unstable partner preference is established, and the 22 h time point is when pair bonding becomes stable (i.e., after overnight mating) (DeVries & Carter, 1999; Williams et al., 1992). It is important to note that the opaque divider in the acclimation period prevented physical interactions, but it is possible that animal pairs may have detected each other through olfactory or auditory cues. The experiment was run in six testing blocks. Each block was composed of eight testing days spread across two weeks, with two adjacent testing days per cohabitation time point (0, 2.5, 6, 22 h). Two pair sets were tested in a day, either male siblings and female siblings or two mating pairs. The order of partner type (adjacent days per time point) and order of cohabitation timepoints (four per block) were counterbalanced across blocks. Individuals were randomly assigned to the partner type condition.

There were 12 sibling pairs and 12 mating pairs tested for each cohabitation time point, for a total of 96 female and 96 male study subjects. Of this total, 190 voles were used for behavioral analyses, and 189 voles were used for IEG analyses. A mating pair from the 0 h timepoint group lacked behavioral data due to a camera malfunction (audio was unaffected).

Three brain samples were not used in IEG analyses: 1 male sibling from the 6 h timepoint group, 1 female mate from the 0 h time point, 1 female mate from the 6 h time point). The first sample was not stained for c-Fos due to major issues with the perfusion. The latter two samples were identified as outliers by a Rosner’s test (EnvStats R package (Millard, 2013)); their whole-brain c-Fos counts were higher than the rest of the samples (R = 4.61 and 5.504, P < 0.05).

### Estrus induction and animal selection

Voles were housed in new home cages, isolated from their sibling cage mates, 4-5 days prior to testing. During this isolation period, all females were induced into estrus with daily 0.1 ml subcutaneous injections of 2 μg estradiol benzoate dissolved in sesame oil (Amadei et al., 2017; Carter et al., 1988). Voles were screened for mating capacity on the 3rd day of isolation. This screening involved a brief exposure (< 10 min) to an opposite-sex vole until the first mount. Voles that did not show a mating attempt in the first exposure were retested with a different animal. We used this mating assay to restrict study subjects to voles that showed lordosis (females) or mounting behavior (males). By selecting voles who showed sexual behavior, we could control the estrus state and timing of mating across the 0, 2.5, 6 and 22 h study groups. This selection process also ensured that animals assigned to the same-sex sibling pair and opposite-sex mating pair groups had similar sexual motivation and experience.

### Behavioral procedures

Behavioral testing began in the morning after lights-on between 8:00 and 10:15 (mean = 8:44 am). Subjects were fitted with colored collars (pipe cleaner attached to a miniature cable tie) for identification and automated video tracking. Then, subjects were placed on opposite sides of a custom made acrylic testing arena (12” x 24” x 12”), separated by an opaque divider, for a 2 h acclimation period. The testing arenas were outfitted with fresh bedding and ad libitum access to chow and water and were housed within an enclosed experiment box (42” x 34” x 24”) constructed from expanded PVC. These experiment boxes had controlled white:red lighting (Phillips Hue light strips) on the same photoperiod cycle as the colony room. For the 0 h time point subjects only underwent the acclimation period. Otherwise, the divider was removed at the end of acclimation so that subjects could interact freely for 2.5, 6 or 22 h. After this acclimation (0 h group) or cohabitation (2.5, 6, 22 h groups), subjects were promptly removed from the arena and perfused. Arenas were cleaned with 70% EtOH between tests.

Video and audio data were recorded from Basler Ace-IMX174 cameras (1920×1200, 2 MP resolution, Basler AG, PA) and Ultramic 364K BLE microphones (Dodotronic, Italy) suspended above each testing arena. Cameras were outfitted with a 16 mm lens and 46mm linear polarizer. Video data were recorded with Pylon viewer (v 5.1.0.12681, Basler AG, PA) at 25 fps with white balance set to ∼1.93 and exposure levels between 15 and 25 ms. Audio was recorded with SeaPro2 software (v 2.0j, CIBRA) at a 192 kHz sampling rate and saved in 30 min WAV files (starting on the hour and every half hour). With this set up, ultrasonic vocalizations (USVs) could be detected per pair, but not localized to specific individuals. Recording equipment was connected directly to a Windows 10 PC. The start of both video and audio recordings were timestamped with PC system time, which enabled the synchronization of video-audio data on a second-by-second timescale.

### Video and audio processing

Behaviors were measured during observation windows that corresponded to peaks in IEG expression (Kim et al., 2015; Renier et al., 2016), specifically 60-120 min before each pair’s perfusion (i.e., midpoint between exsanguination times of each vole pair). Both automated and manual methods were used to characterize vole behavior.

Automated scoring of movement and proximity behaviors was done with Ethovision (v 10.1). Individual collars were tracked during white light periods, with detection settings set for each video and optimized for collar color (blue / green, mean ± sd: huemin = 97.0 ± 1.4 / 63.1 ± 2.5, huemax = 113.0 ± 1.4 / 79.2 ± 2.6, saturationmin = 113.3 ± 11.8 / 85.4 ± 10.5, saturationmax = 255.0 ± 0.2 / 254.9 ± 0.7, brightnessmin = 84.1 ± 10.5 / 70.2 ± 8.9, brightnessmax = 254.0 ± 4.6 / 254.7 ± 1.6, marker size = 50). Body area was recorded across both white light and red light periods with the grayscaling method (white / red light: detectionmin = 0, detectionmax = 108.7 ± 3.3 / 5.96 ± 0.8, pixel sizemin = 3,000, pixel sizemax = 125,000, contour erosion = 1, contour dilation = 1). The largest body area was recorded when animals were separated. Overall activity (% pixel change between frames) also was recorded for white light and red light periods separately (white / red light: threshold = 1, background = 10, compression = “on” for all 24 overnight videos / “on” for 18 overnight videos). Individual tracking, body area, and activity were recorded at 12.5 samples/s, down sampled to 1/s, and used to compute an array of behavioral measures across entire recording sessions and the 1 h IEG induction windows (Figure 2—figure supplement 1A,B and Supplementary File 3).

Manual scoring of mating, investigative, grooming, and contact behaviors was done with BORIS (v 7.9.6) (Friard & Gamba, 2016). Two trained observers, who were blind to study animals’ sex and partner condition, independently labelled social and non-social behaviors in cohabitation videos (2.5, 6, 22 h groups) during the 1 h IEG induction windows (Supplementary File 3). Self-grooming during acclimation videos (0 h group) was labelled by one of the trained observers. Inter-observer reliability was assessed with Pearson correlations. For dyadic behaviors (e.g., mounting, huddling), reliability was assessed across vole pairs. For individual behaviors (e.g., anogenital sniffing, self-grooming), reliability was assessed separately for individuals with blue collars and green collars. All social behaviors had high agreement between independent observers (Pearson correlations, r > 0.90; Supplementary File 3).

Ultrasonic vocalizations (USVs) were detected with DeepSqueak (v 2.6.1 in MATLAB 2019a) (Coffey et al., 2019). To initially label USVs, the “AllShortCalls” network was used along with the following detection settings: frequency range of 20-90 kHz, overlap of 0.1s, 5s chunk length, and a high precision recall. Then, a trained observer revisited all USVs labels during the 1h behavior observation windows. Using the DeepSqueak interface, the observer removed labels of background noise (e.g., water bottle sounds) and adjusted USV label boundaries to exclude noise. This trained observer was blind to the experiment condition (siblings vs mates), caller sex, and the behavioral context. The start and stop times of all USV labels were exported and the times were adjusted to align with the video frame timestamps.

### Statistical analyses

Absolute and relative volumes of brain regions were compared across species (mouse vs. vole) and sexes (male voles vs. female voles) with negative binomial regression models. Relative volumes were computed as the ratio of region volumes to whole brain volumes. To control for false discovery rate, p-values were corrected for multiple comparisons to q values (Hochberg, 2015).

All statistical analyses for behavioral experiments were carried out in R (v 3.5.3) (R Development Core Team, 2016). Welch two-sample t-tests were used to compare behavioral measures between mate pairs and sibling controls. Paired t-tests were used to compare behavioral measures between female mates and male mates. T-tests were run for all timepoints combined and for specific timepoints. Significance values were adjusted for multiple comparisons using the FDR method, with an alpha threshold of q=0.05. Pearson correlations were used to make associations between behavioral measures and ROI c-Fos+ cell counts.

To identify brain regions that are sensitive to pairing status, we used a model comparison approach. For each voxel or ROI, two general linear regression models (“GLMs”; a quasi-poisson link function was used to model over-dispersed count data) were used to predict IEG c-Fos expression across all individuals (n=189) (Figure 3, Video 2). These models differed in whether the partner condition (sibling vs. mate) was a part of the formula. The “null” model (Equation 1) included main effects of sex [S], time point [T], and experiment block [B]. The hypothesized “bonding” model (Equation 2) included the same predictor terms in addition to a main effect of partner type [P] and interactions between partner with sex or timepoint. In these models, sex was a categorical factor (males vs females). Time point was an ordinal variable from 1 to 4 (0, 2.5, 6, 22h) and was included as a polynomial term to account for both linear and quadratic (i.e., 0/22h vs 2.5/6h) effects. Block was an ordinal variable from 1 to 6. Partner was a categorical variable (mates vs siblings). This model comparison allowed for the identification of voxels/ROIs where c-Fos expression variation was explained better when accounting for the partner condition.

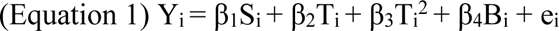

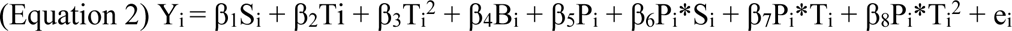

The null and bonding GLMs were compared with an ANOVA test. For the ROI-level analysis, a Monte Carlo approach (10,000 random shuffles of the data) was used to determine a null distribution for the ANOVA F-statistics and compute p-values. For the voxel- and ROI-level analyses, the FDR method was used to correct for multiple tests, with an alpha threshold of q=0.1.

We ran an additional model comparison to specifically test for sex differences in c-Fos expression (Video 2). In this analysis, we compared the full version of the model formula (Equation 2) to a reduced version of the model (Equation 3). The reduced model excludes a sex by partner interaction term. The FDR method was used again to correct for multiple tests across all voxels and all ROIs, with an alpha threshold of q=0.1.

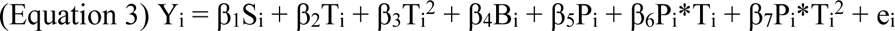

ROIs were chosen for subsequent analyses if they were significant in the Equation 1 vs. Equation 2 model comparison and were mutually exclusive with one another. To select mutually exclusive ROIs, we constructed an structural hierarchy from the ARA. For any significant ROIs that overlapped anatomically, we iteratively chose the ROI with the higher ANOVA F-statistic. This method excluded larger brain regions in lieu of smaller and more localized ROIs.

Significant “chosen” ROIs were assigned to groups by using hierarchical clustering with the ward D2 method with a Euclidean distance matrix extracted from c-Fos cell counts (Murtagh & Legendre, 2014; R Development Core Team, 2016). The resulting tree was cut so that ROIs were grouped into anatomically similar clusters. We used multi-dimensional scaling (MDS) to further interpret the degree of similarity between chosen ROIs and their clustering groups based on the Euclidean distance matrix. The MDS method is a form of non-linear dimensionality reduction to visualize similarity in Cartesian space (Cox & Cox, 2008).

To confirm whether chosen ROI clusters represented structural circuits, we compared cluster assignments to published data on structural connectivity in the mouse brain (Knox et al., 2019). First, we refined a matrix of ROI-ROI ipsilateral normalized connection densities to align with our list of chosen ROIs. Some of our chosen ROIs did not align with this matrix because they represented subregions of the ROIs in the matrix. In those cases, we used data from the next inclusive ROI that was available (e.g., BST data used for BSTpr). Then, for each cluster, we found the mean normalized connection density between the regions. We excluded data from the matrix diagonals (e.g., BST to BST) to emphasize connections between, rather than within, regions. We took an average of these cluster densities values to capture the overall connection density based on our cluster assignments. A permutation test was used to assess whether this connection density was higher than expected by chance. The rows of the connectivity matrix (rows = origin ROIs, columns = target ROIs) were randomly shuffled prior to computing the average cluster density, and this was done 10,000 times to construct a null distribution. This null distribution was then compared to the observed density to estimate its probability.

We used canonical-correlation analysis (CCA) was used to investigate the relationships between behavioral variables and IEG induction patterns in chosen ROIs. CCA is an unsupervised method that finds linear combinations of two variable sets with the strongest correlation (González et al., 2008; R Development Core Team, 2016; Wang et al., 2020). This approach enabled us to isolate discrete canonical correlates (CC), where each individual animal is assigned scores for each variable set per CC factor. We used correlations between individual behavior/ROI measures and their CC scores to interpret CC factors and identify specific behaviors and ROIs with the strongest associations. Wilk’s lambda test statistic was used to confirm which CC factors represented a significant association between the two variable sets.

## Data and code availability

Source data and source code, along with the prairie vole reference brain and atlas, are available on Figshare (DOI: 10.6084/m9.figshare.21375666). Raw behavioral and neural data will be made available upon reasonable request.

## Acknowledgments

We thank animal resource staff at the University of Texas at Austin and Cold Spring Harbor Laboratory for their assistance with animal care and husbandry. Tracy Burkhard helped with animal perfusions and tissue collection, and D. Lama and M. Klein assisted with behavior scoring. We thank Janelle Collins and other staff at Certerra Inc. for their assistance with immunohistochemistry, whole-brain imaging and preliminary analyses, and we thank Kith Pradhan for advice on customizing the voxel and ROI analysis pipeline. Rhonda Drewes and Jason Palmer assisted with alignment of the reference atlas to the prairie vole brain and with imaging. Jessica Tollkuhn provided additional prairie voles for species comparisons and immunohistochemistry. Finally, we thank Drs. Hans Hofmann. Adam Kepecs, Micheal Long, and Michael Ryan for their feedback on earlier versions of this manuscript. This research was funded by grants from the National Institutes of Health: R01MH115267 (S.M.P., P.O.) and K99MH126164 (M.L.G.).

## Author contributions

All authors conceptualized the research. M.L.G. and R.M.C. performed the research and analyses. S.M.P. and P.O. supervised the research. M.L.G. and S.M.P. wrote the paper. All authors reviewed and edited the paper.

## Figure supplements

**Figure 1—figure supplement 1.**
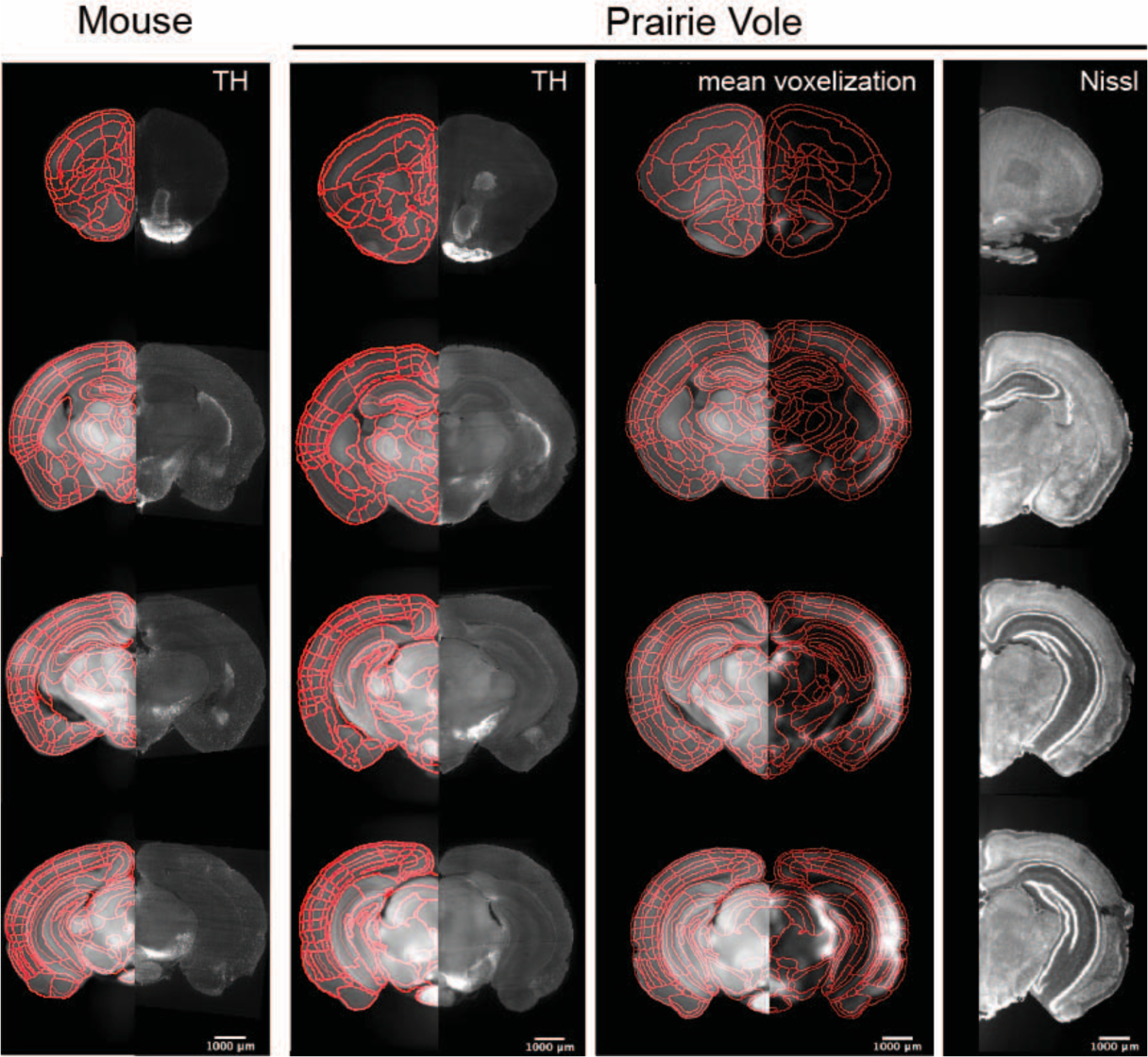
Validation of the prairie vole reference atlas. Mouse coronal sections (left column) are composed of the mouse reference brain overlaid with atlas boundaries in red on the left and, in the same sections, of tyrosine hydroxylase (TH) immunolabeling on the right. Prairie vole coronal sections are in the right three columns. In the first vole column, coronal brain sections are of the prairie vole reference brain and atlas boundaries in red on the left with TH immunolabeling on the right. In the second vole column, coronal sections are of the prairie vole reference brain and c-Fos+ mean voxelization overlay with atlas boundaries in red. In the third vole column, coronal sections of prairie vole NeuroTrace staining are registered onto the prairie vole reference brain.

**Figure 2—figure supplement 1.**
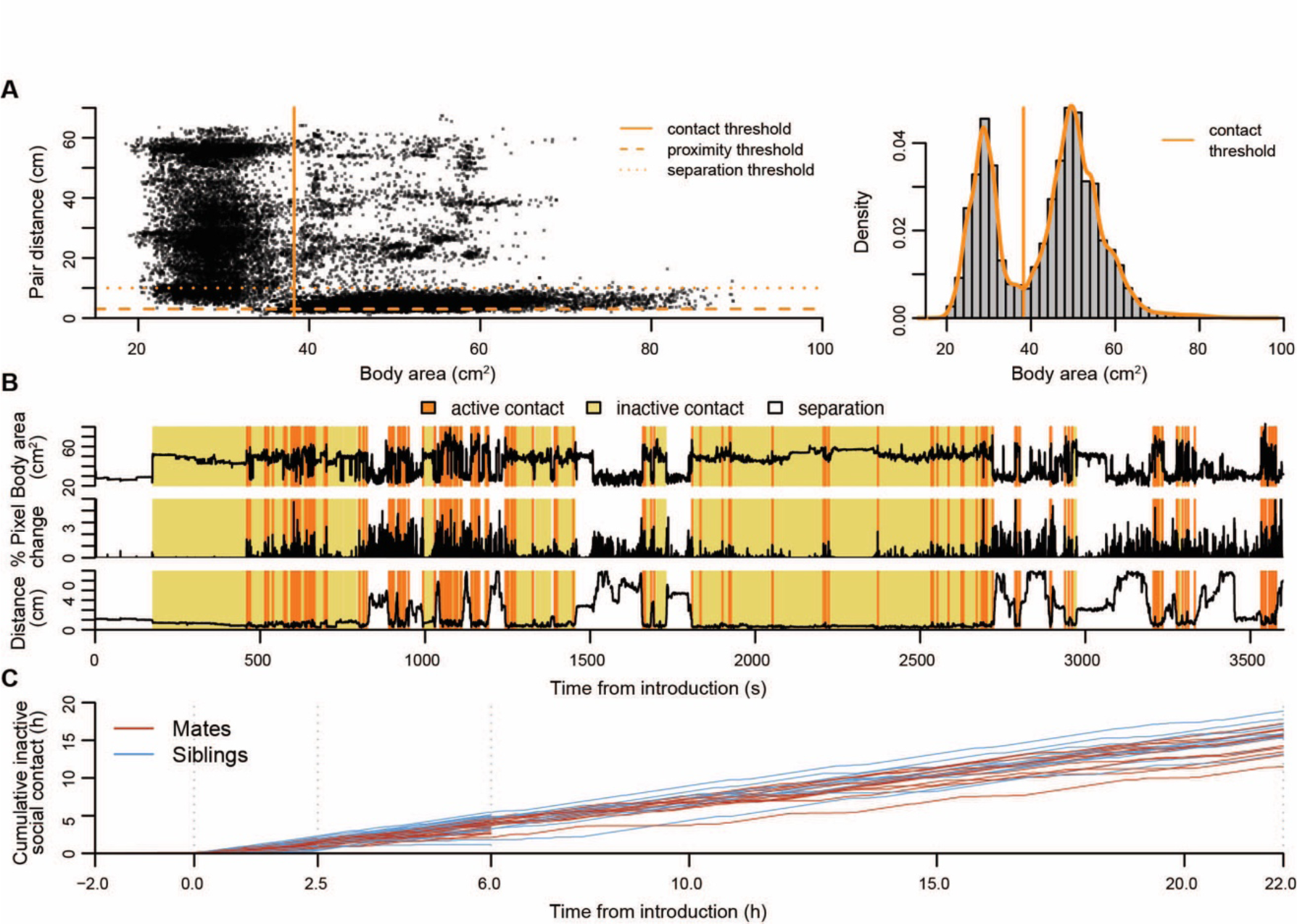
Automated tracking of social behavior states. **(A)** Long-term tracking of social states was informed by automated measures of the largest body area, pair activity, and pair distance. On the left, body area is plotted against dyad distance from an exemplar 24h mate pair video (10% randomly selected frames from 12 hours of white light). On the right, the density curve for body area reveals a basic threshold for when animals are separated or in physical contact. **(B)** Traces of body area, video activity, and pair distance are plot for the first hour of interaction of the exemplar mate pair, along with automated assignments of behavioral states. **(C)** Cumulative time spent in inactive social contact for up to 22h of cohabitation in mate pairs (red lines) and sibling pairs (blue lines).

**Figure 2—figure supplement 2.**
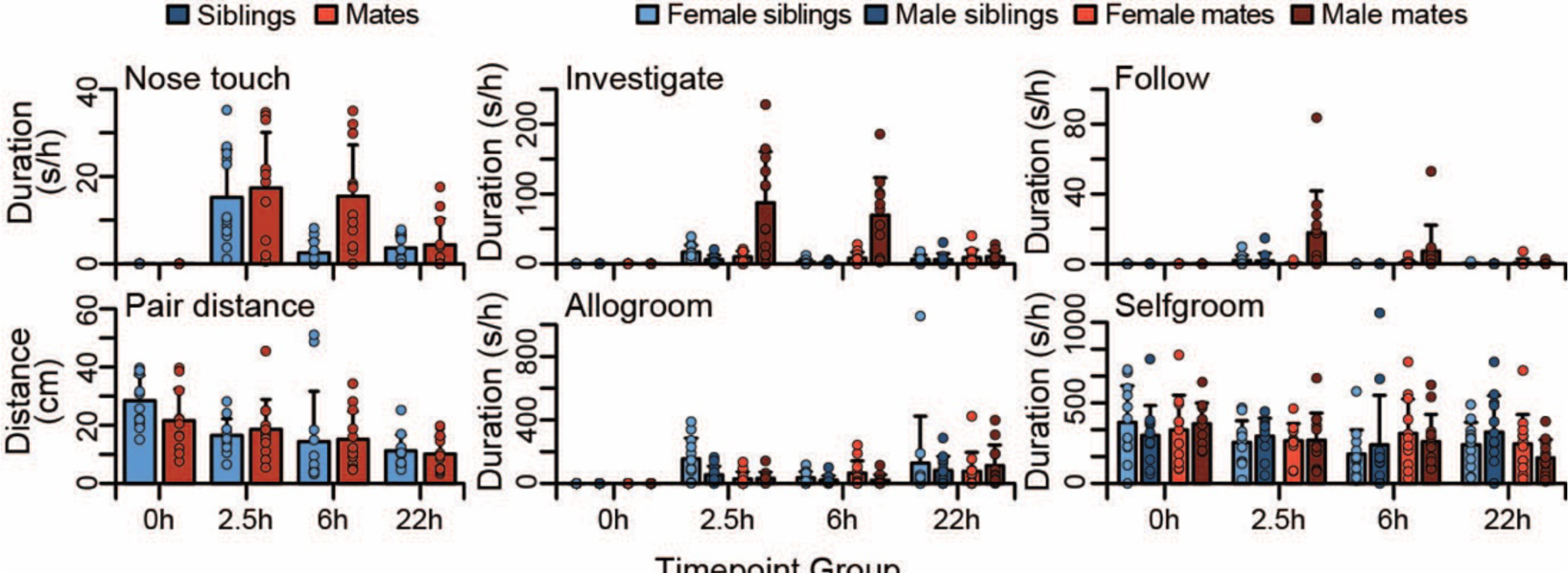
Time course of social behaviors during pairing. Group differences (mean ± sd) are shown for appetitive behaviors including nose-to-nose touching, anogenital investigation, and close follows. Group differences (mean ± sd) are shown for proximity and grooming behaviors including pair distance, allogrooming, and selfgrooming. Mate pairs are in red and sibling pairs in blue (female in lighter hues, males in darker hues). T-tests were used to compare mates and siblings, and paired t-tests were used to compare female and male mates.

**Figure 2—figure supplement 3.**
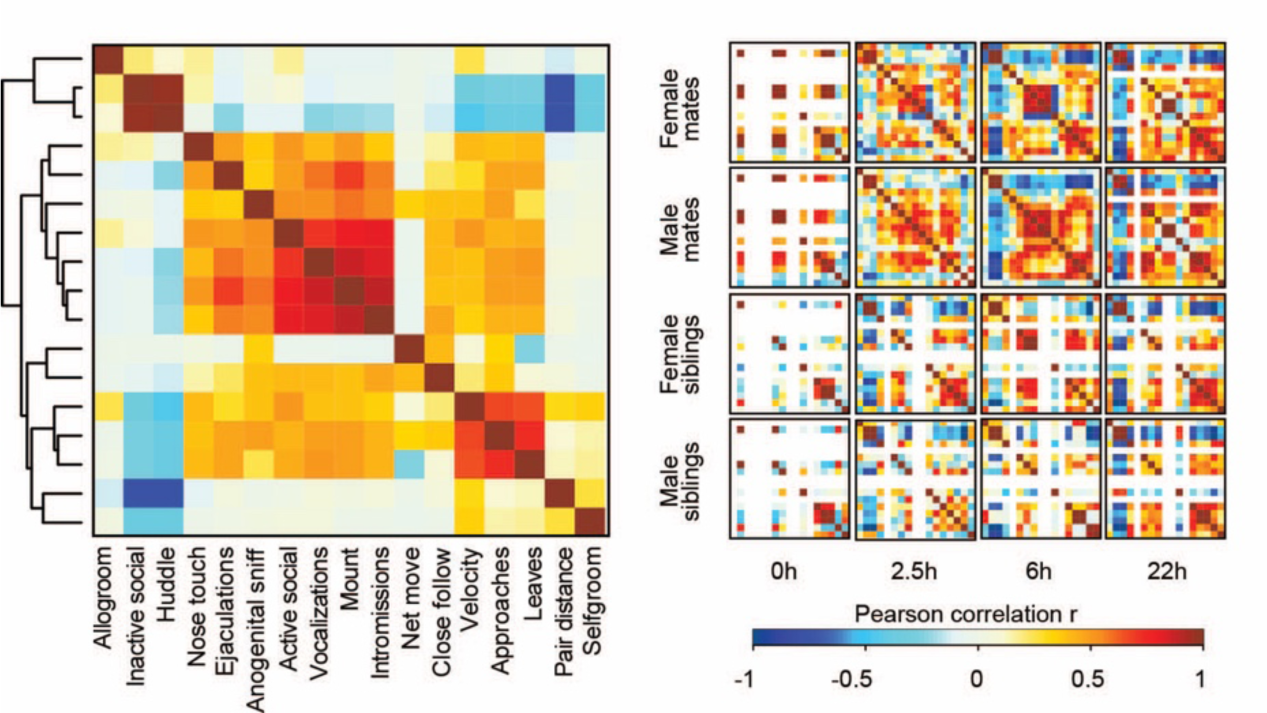
Associations between behavioral states and types of social interaction. Hierarchical clustering of behavioral measures from Pearson correlations groups behavioral states and interactions into three main clusters involving close contact (e.g., huddling and allogrooming), mating (e.g., mounts, vocalizations), and appetitive behavior (e.g., approaches and follows). On the right, Pearson correlations are shown among behaviors in subsets based on partner type, sex, and timepoint. Warm and cool colors indicate positive and negative correlation coefficients, respectively. White indicates behaviors with no variation, meaning that coefficients were not computed (e.g., siblings did not exhibit mating behaviors).

**Figure 3—figure supplement 1.**
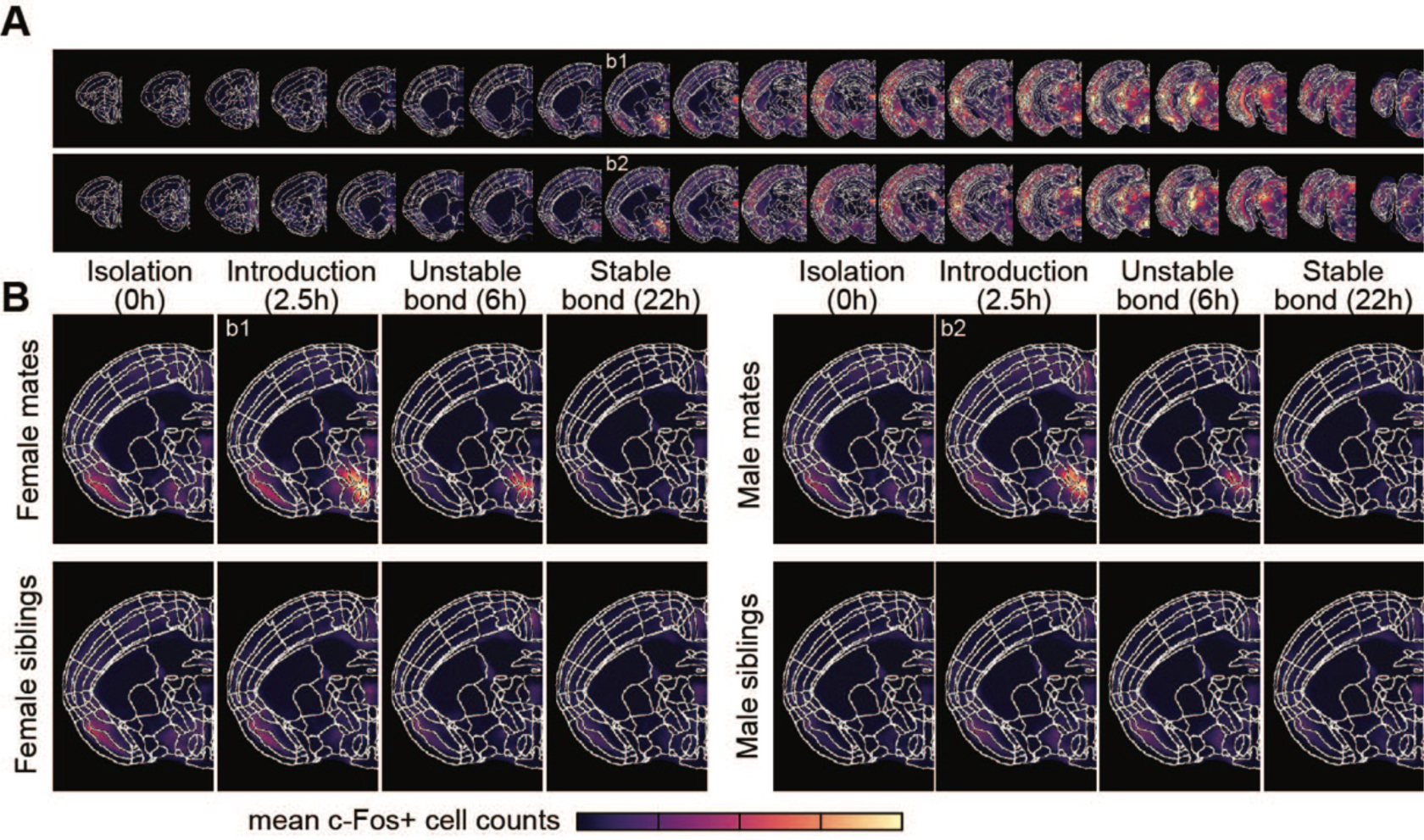
Brain-wide patterns of immediate early gene activation during pairing. **(A)** Coronal cross-sections (rostral to caudal) from female (top) and male (bottom) mating pairs are shown for the 2.5h timepoint group, with brightness corresponding to the average voxel c-Fos+ cell counts. **(B)** A representative coronal slice that includes posterior BST is shown for mate pairs and siblings and separated by timepoint and sex. Brightness corresponds to average voxel c-Fos+ cell counts per group.

**Figure 3—figure supplement 2.**
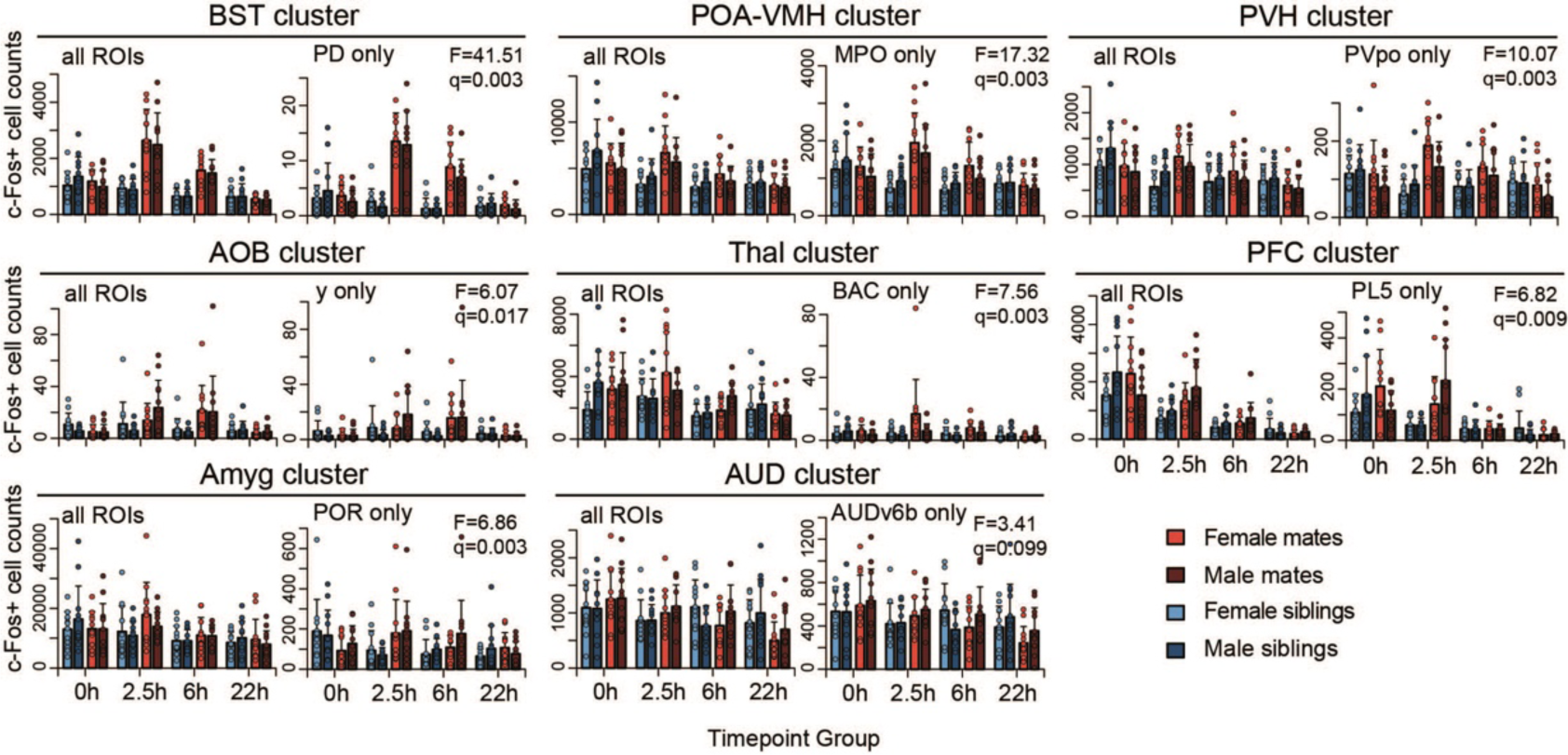
Patterns of immediate early gene expression in brain region clusters. Counts of c-Fos+ cells are shown for ROIs associated with pair bond development, organized by hierarchal cluster. For each cluster, the total counts are shown on the right and the most significant ROI (highest F-statistic) is shown on the right. Counts are summarized by timepoint, partner type and sex (mean ± sd), and overlaid with counts from individual animals. Mate pairs are in red and sibling pairs in blue (female in lighter hues, males in darker hues). Test statistics are from an ANOVA to compare null and hypothesized general linear models. FDR q-values are from permutation tests to assess the likelihood of the observed test statistic as compared to a distribution of test statistics obtained from shuffled data of 10,000 permutations.

**Figure 3—figure supplement 3.**
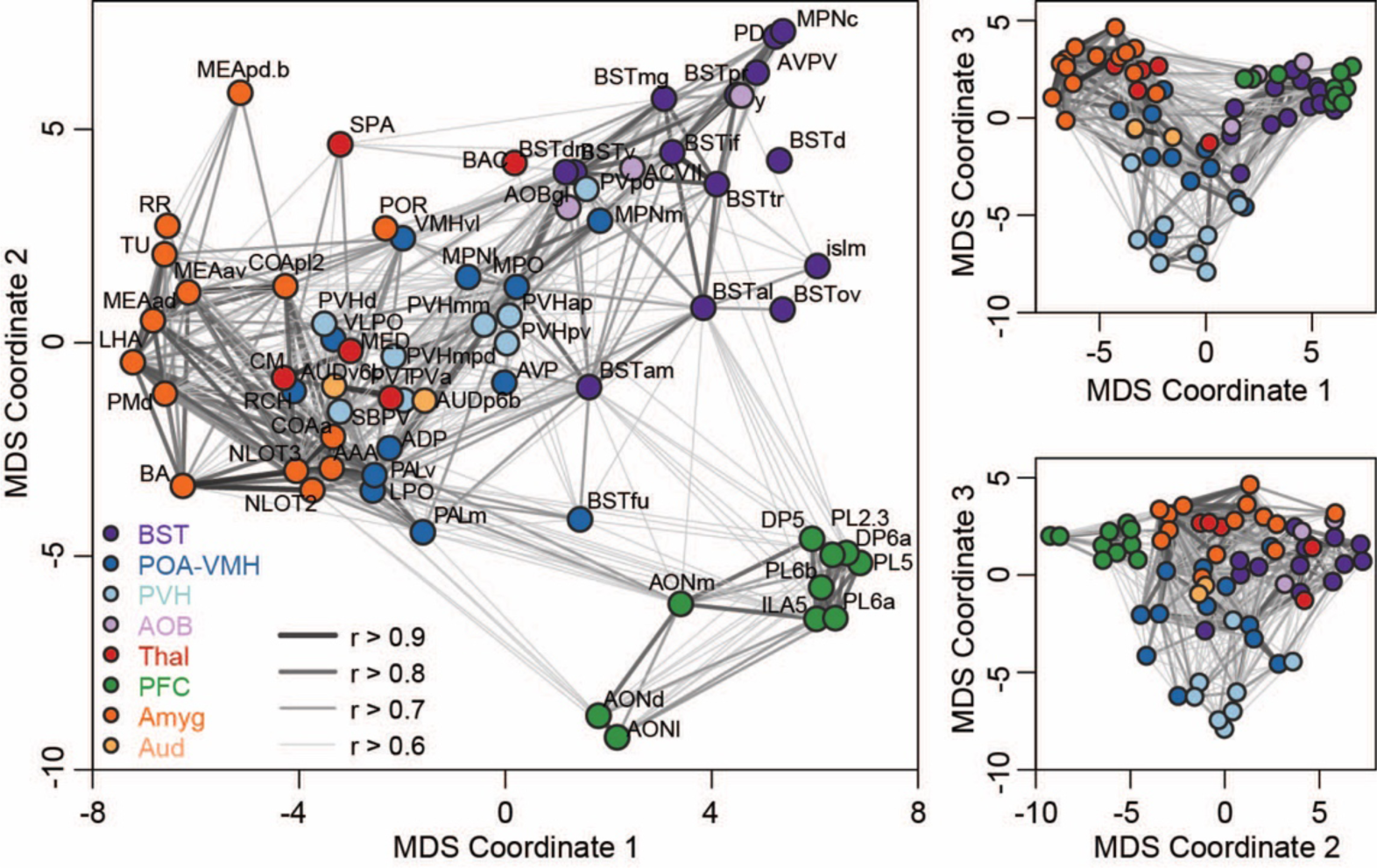
Multi-dimensional structure of brain-wide correlation patterns. Representations of the first three dimensions of a multi-dimensional scaling (MDS) coordinate space based on Pearson correlations between c-Fos+ cell counts in brain regions (ROIs) associated with bonding. Each symbol represents an ROI and is colored based on cluster assignment. Darkness and thickness of connecting lines reflect correlation coefficients.

**Figure 3—figure supplement 4.**
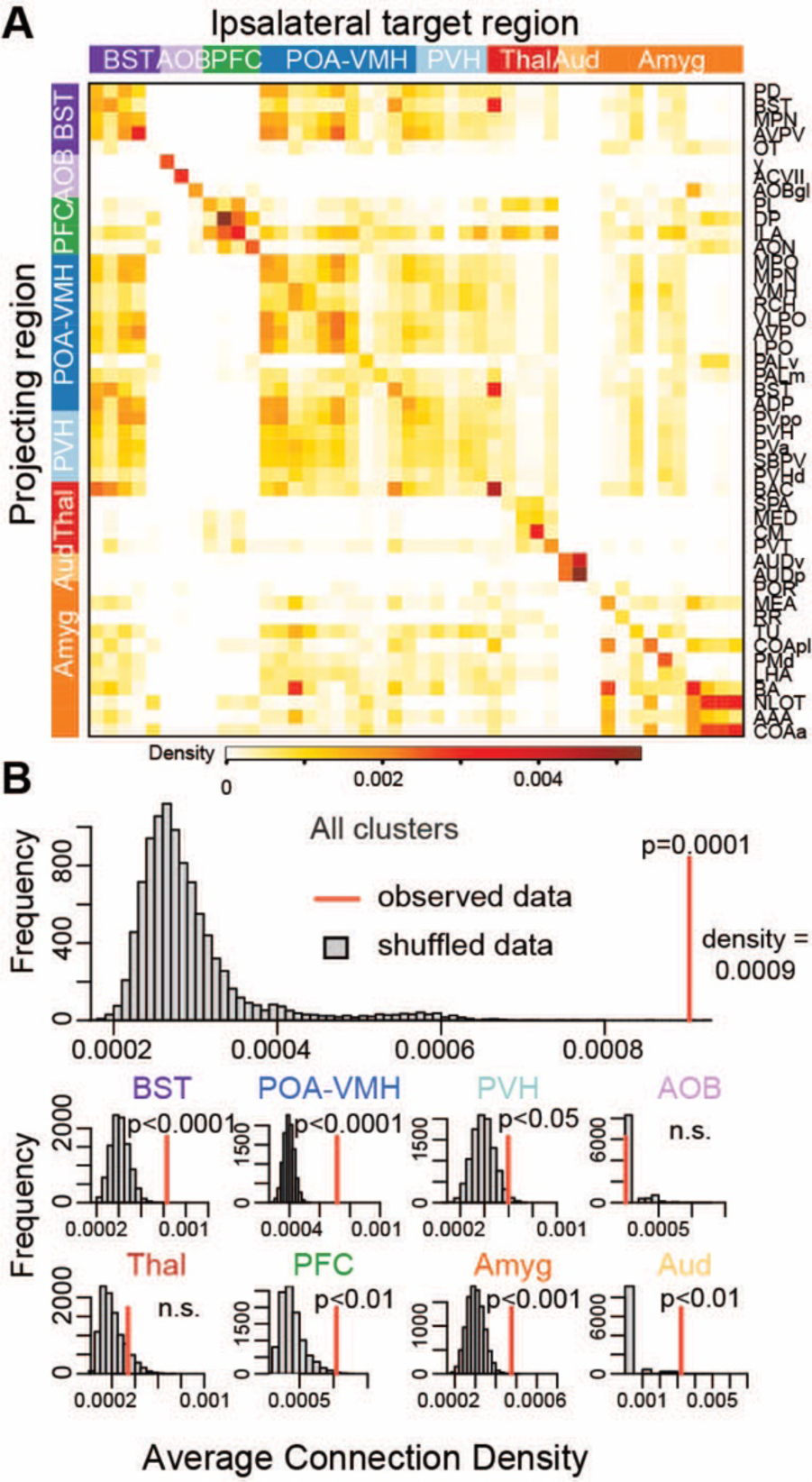
Anatomical connectivity in brain regions associated with pairing. **(A)** This heatmap shows normalized connection densities (Knox et al., 2019) from projecting to ipsilateral target ROIs in the mouse brain. These ROIs are the same as, or larger divisions that contain, the ROIs found to be associated with pair bond development in our analysis. **(B)** The top histogram shows results of a permutation test to assess whether hierarchical clusters of chosen ROIs in our analysis mirror underlying anatomical connections. Observed connectivity reflects the overall average of cluster means, which is compared to averages from 10,000 iterations of shuffled data (i.e., target regions shuffled for each projecting region). The bottom histograms show the results of permutation tests for the mean connectivity within each cluster of ROIs.

**Figure 4—figure supplement 1.**
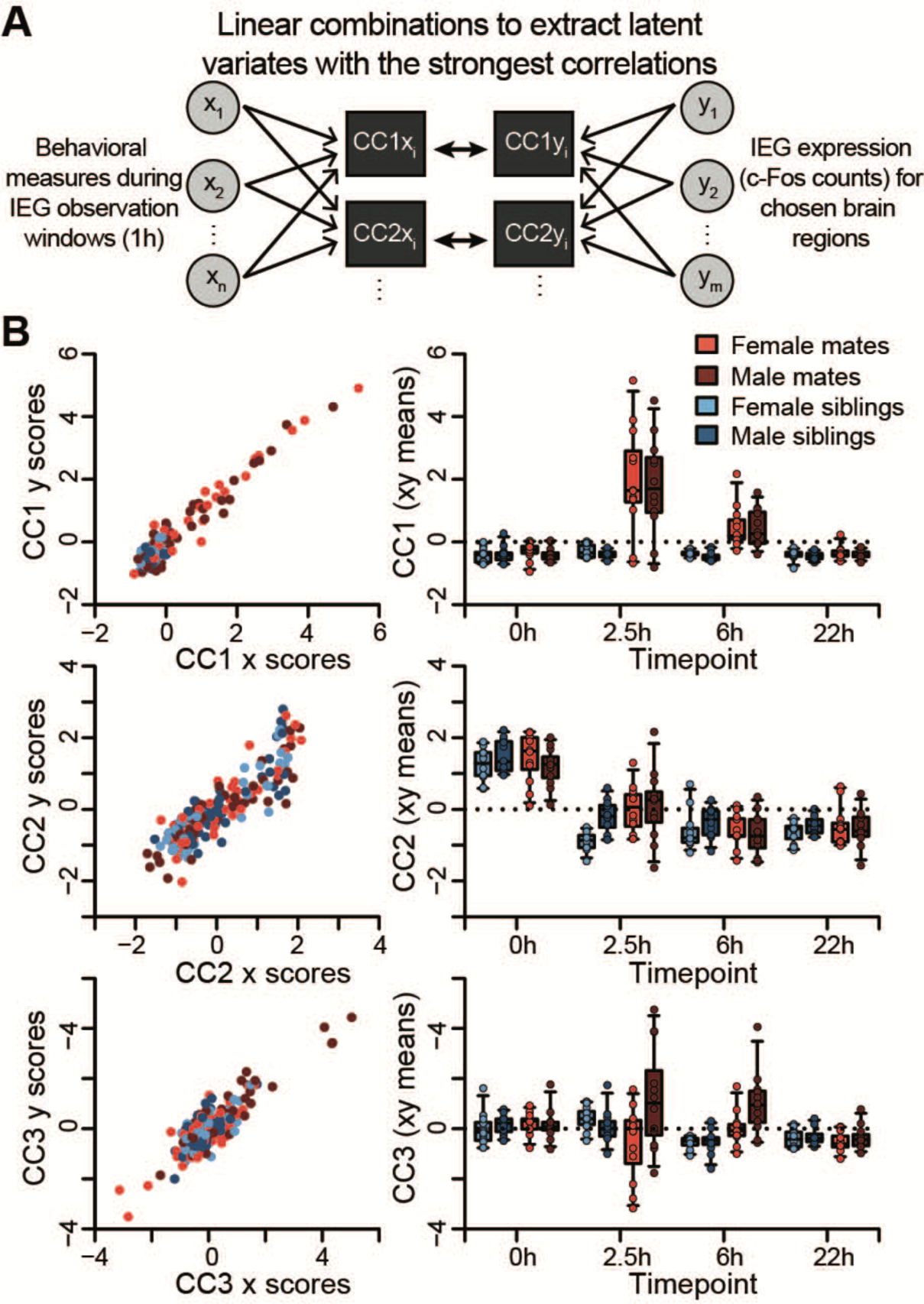
Dimensions of cross-covariance in immediate early gene expression and social behavior. **(A)** Schematic of canonical correlation analysis (CCA), where two sets of variables (x and y) are combined linearly to extract latent variates with the strongest correlations (CC1 stronger than CC2, and so on). This analysis outputs x and y scores for each latent variate for each animal subject in the dataset. In this dataset, x scores represent linear combinations of behavioral measures, and y scores represent linear combinations of ROI c-Fos counts. **(B)** On the left, the relationship between x and y scores is shown for all study animals for the first three variates. On the right, the means of x and y scores for the first three variates are compared across partner type, sex, and timepoint (boxplots = median, 25%-75% quartile, and 95% confidence intervals).

**Figure 4—figure supplement 2.**
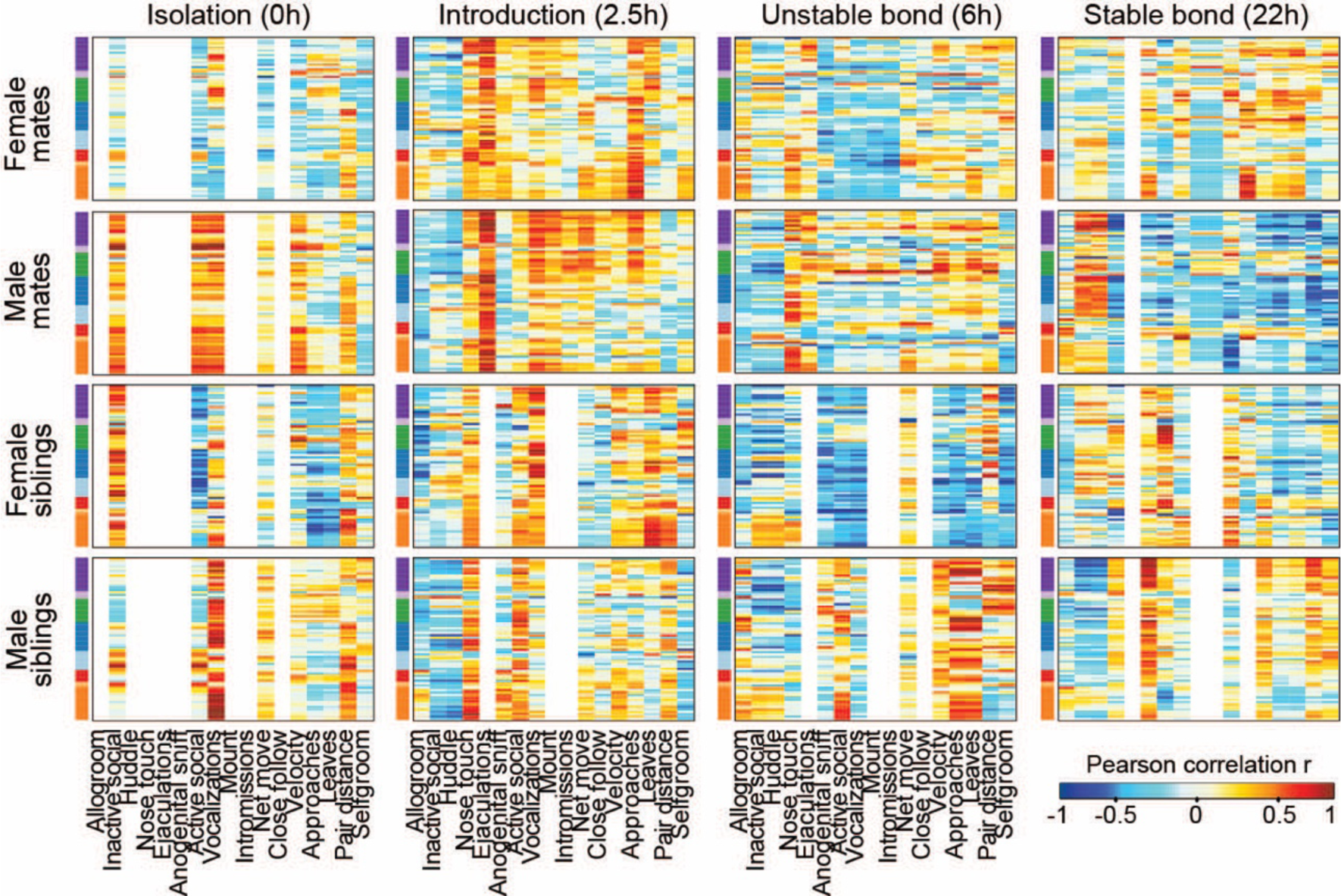
Intra-pair similarity in immediate early gene expression in pairing-associated brain regions. Heatmaps represent Pearson correlation coefficients, with brain regions on the y-axis (grouped into hierarchical clusters, see Figure 3 for ROI and cluster labels) and behavioral outcomes (during 1h observation windows) on the x-axis. Correlation heatmaps are split by partner type, sex and timepoint. Warm and cool colors indicate positive and negative correlation coefficients, respectively. White indicates behaviors with no variation, meaning that coefficients were not computed (e.g., siblings did not exhibit mating behaviors).

**Figure 4—figure supplement 3.**
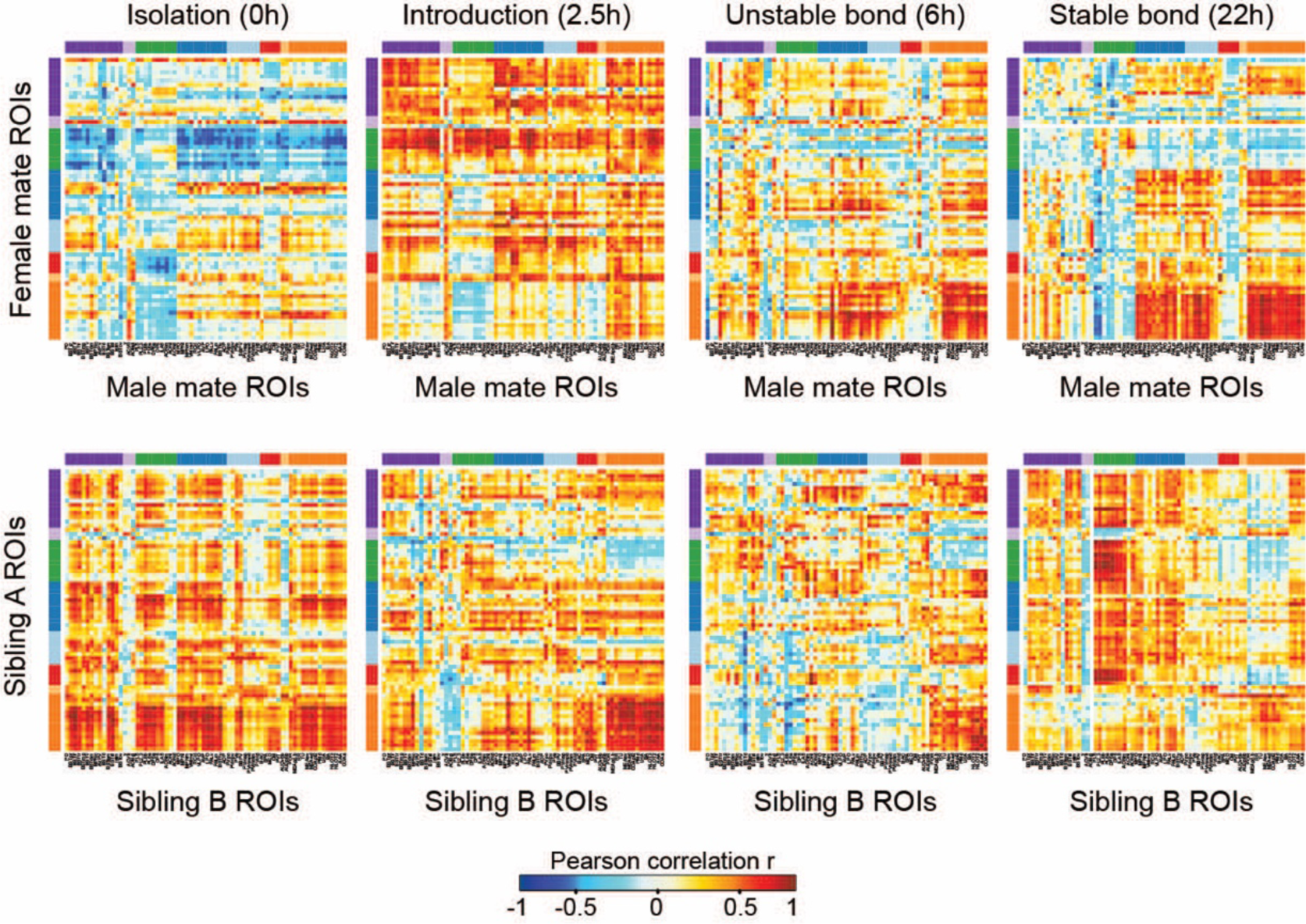
Patterns of association between immediate early gene expression and behavior. On the top, heatmaps represent Pearson correlation coefficients between selected brain regions (ROIs) in mating pairs, with female data on the y-axis and male data on the x-axis. On the bottom, heatmaps represent Pearson correlation coefficients between selected ROIs in sibling dyads, where sibling A data are from animals placed in the left side of chamber during acclimation, and sibling B data are from animals placed on the right side. ROIs are grouped by hierarchical clustering (see Figure 3). Warm and cool colors indicate positive and negative correlation coefficients, respectively.

## Supplementary and Rich Media files

**Supplementary File 1. Statistical comparisons of absolute and relative brain area volumes between prairie voles and mice.** Absolute and relative area volumes (normalized to total brain volume) are compared between prairie voles and male mice. Absolute brain area volumes are all statistically significant bigger in the prairie vole compared to the mouse, but none the ratios of area volume (relative to the whole brain) were statistically different between species. Mean, sd, p-values, and FDR correction of the p-values are presented for each region.

**Supplementary File 2. Statistical comparisons of absolute and relative brain area volumes between male and female prairie voles.** Absolute and relative area volumes (normalized to total brain volume) are compared between male and female prairie voles. No statistical differences were found in these sex comparisons. Mean, sd, p-values, and FDR correction of the p-values are presented for each region.

**Supplementary File 3. Prairie vole behavior ethogram.** Descriptions of automated, semi-automated, and manually scored behavioral measures.

**Supplementary File 4. Statistical results from comparisons of general linear models across brain regions**. Null general linear models (GLMs) and hypothesized (i.e., “bonding”) GLMs were compared with ANOVA tests for each region of interest. The ANOVA F-statistics are reported alongside p-values computed from permutation tests (10,000 shuffles). The corresponding q-values were computed with the FDR method to correct for multiple tests.

**Video 1. Prairie vole reference brain.** The prairie vole reference brain from LSFM imaging is shown from coronal cross-sections (rostral to caudal) and in a 3D view. Then, coronal cross-sections are shown of the prairie vole atlas (left) alongside whole-brain staining for somatostatin (SST, middle) and fluorescent Nissl (NeuroTrace, right).

**Video 2. Patterns of whole-brain immediate early gene expression during pairing.** First, coronal cross-sections of average IEG induction patterns are shown for female (first video) and male (second video) prairie voles across each of the 16 experiment groups split by partner type (mates-top, siblings-bottom) and timepoint (0, 2.5, 6, 22 hours from left to right). Brightness corresponds to average voxel c-Fos+ cell counts. These videos are associated with Figure 3— figure supplement 1; see these panels for the color key. The induction patterns are overlaid with prairie vole atlas boundaries. Next, coronal cross-sections are shown with the results of model comparisons to identify brain areas associated with bonding. This video is associated with Figure 3A (see this panel for the color key). The reference is overlaid with colored voxels that represent a difference (FDR q < 0.1) between hypothesized general linear model that included partner type and a null model that did not. Warm colors indicate voxels with higher c-Fos+ counts in mate pairs (test statistic for partner type > 0), and cooler colors indicate voxels with higher c-Fos+ cell counts in siblings (test statistic for partner type < 0). Finally, coronal cross-sections are shown with the results of model comparisons to identify brain areas associated with sex differences. The reference is overlaid with colored voxels that represent a difference (FDR q < 0.1) between hypothesized general linear model that included a partner type x sex interaction term and a null model that did not. Colored voxels indicate regions where there was an interaction effect, with warm colors indicating a positive interaction term and cool colors indicating a negative interaction term (see Figure 3A color key).

